# Phylogenetic profiling and cellular analyses of ARL16 reveal roles in traffic of IFT140 and INPP5E

**DOI:** 10.1101/2021.10.14.464442

**Authors:** Skylar I. Dewees, Romana Vargová, Katherine R. Hardin, Rachel E. Turn, Saroja Devi, Joshua Linnert, Uwe Wolfrum, Tamara Caspary, Marek Eliáš, Richard A. Kahn

**Author notes:** Corresponding author: Richard A. Kahn,; Department of Biochemistry, Emory University School of Medicine, 1510 Clifton Rd, Atlanta, GA, USA 30322.

## Abstract

The ARF family of regulatory GTPases is ancient, with 16 members predicted to have been present in the last eukaryotic common ancestor. Our phylogenetic profiling of paralogs in diverse species identified four family members whose presence correlates with that of a cilium/flagellum: ARL3, ARL6, ARL13, and ARL16. No prior evidence links ARL16 to cilia or other cell functions, despite its presence throughout eukaryotes. Deletion of ARL16 in MEFs results in decreased ciliogenesis yet increased ciliary length. We also found *Arl16* KO in MEFs to alter ciliary protein content, including loss of ARL13B, ARL3, INPP5E, and the IFT-A core component IFT140. Instead, both INPP5E and IFT140 accumulate at the Golgi in *Arl16* KO lines, while other IFT proteins do not, suggesting a specific defect in traffic from Golgi to cilia. We propose that ARL16 regulates a Golgi-cilia traffic pathway and is required specifically in the export of IFT140 and INPP5E from the Golgi.

**Summary:** Phylogenetic analyses of ARF family GTPases predict that ARL16 is linked to cilia. This was confirmed using MEFs deleted for ARL16, resulting in defects in Golgi to cilium traffic, with accumulation of IFT140 and INPP5E at Golgi.

## INTRODUCTION

ARF family regulatory GTPases are best known for their roles in regulating bi-directional membrane traffic throughout the secretory pathway, including at Golgi and endosomes (Sztul et al., 2019). However, they also regulate a diverse array of other pathways and functions, including at cilia (Fisher et al., 2020), mitochondria, lipid droplets, centrioles, midbodies, and rods and rings (Sztul et al., 2019). To achieve such widespread cellular regulation, mammals express as many as 30 ARF family GTPases, including 6 ARFs, 22 ARF-like (ARL), 2 SARs, and TRIM23. Like all regulatory GTPases, they switch between “inactive” (GDP-bound) and “active” (GTP-bound) conformations in response to the actions of guanine nucleotide exchange factors (GEFs) and GTPase activating proteins (GAPs). With regulatory roles in such diverse and essential cellular processes, ARF family members are implicated in a large number of pathologies including cancers, ciliopathies, hearing and vision impairments, idiopathic pulmonary fibrosis, and other maladies. Despite their importance to cell biology and human health, we lack substantial mechanistic details of their actions (Sztul et al., 2019). Indeed, some of the core family members remain virtually unstudied.

We recently reported the results of a taxonomically broad phylogenetic analysis of the ARF family in eukaryotes (Vargova et al., 2021). This study defines the set of family members ancestral for eukaryotes and describes 16 paralogs putatively present in the last eukaryotic common ancestor. We used our large phylogenetic dataset to look for the retention or loss of family members that are correlated with retention or loss of the cilium. Strikingly, as elaborated below, the phyletic pattern of ARL16 genes provided a hypothesis that its cellular function is relevant to cilia. A link between ARL16 and cilia also emerged in a CRISPR screen for regulators of the Hedgehog signalling pathway (Breslow et al., 2018). ARL16 was previously reported to be cytosolic and inhibit the function of the RIG-I protein in defense against RNA viruses (Yang et al., 2011). Also, two high-throughput datasets identified potential interacting partners of ARL16, including PDE6D (aka PDE6δ or PrBP/delta) and GM130, providing additional clues to its location and functions (Luck et al., 2020; Rolland et al., 2014). Thus, despite some fragmentary data on ARL16 from several systems there has been no systematic approach aimed at identifying fundamental actions of ARL16 in cells, or more specifically in cilia.

The primary cilium is a sensory structure, responding to extracellular signals with cellular responses through complex intracellular signalling networks. Primary cilia are made up of a microtubule-based backbone, the axoneme, which extends from the modified mother centriole, termed the basal body (Satir and Christensen, 2007). The axoneme is covered in the ciliary sheath, which is continuous with the plasma membrane but functionally separated from it by the transition zone, which tightly regulates entry to the ciliary compartment (Stephen et al., 2017). Three paralogs in the ARL family, ARL3, ARL6, and ARL13, play roles in ciliary functions, acting both inside and outside of cilia (Chiang et al., 2004; Fan et al., 2004; Fisher et al., 2020; Gigante and Caspary, 2020; Zhou et al., 2006). ARL3 regulates the delivery of N-myristoylated and prenylated proteins to the cilium by binding cargo transporters UNC119 and PDE6D, respectively (Fansa et al., 2016; Stephen et al., 2017). ARL6/BBS3 regulates the function of the BBSome (a protein complex involved in export of ciliary membrane proteins (Mourão et al., 2014). ARL13, including the extensively studied mammalian paralog ARL13B, is involved in ciliary protein import and export, partly mediated by its activity as a GEF for ARL3 (Gotthardt et al., 2015; Ivanova et al., 2017). It is also well known for its crucial role in vertebrate development via one of the best-known ciliary signaling pathways, Hedgehog (Hh) (Caspary et al., 2007). In this pathway, Hh ligand binds to its receptor, Patched (PTCH) that is enriched on the ciliary membrane. This induces the removal of PTCH from the cilium, allowing the G protein coupled receptor (GPCR) Smoothened (SMO) to enter (Goetz et al., 2009). This leads to cleavage and activation of Gli transcription factors which then translocate to the nucleus, resulting in the transcription of target genes. This pathway is heavily influenced by other ciliary proteins including ARL13B, ARL3, and INPP5E, often through alterations in SMO recruitment or changes in Gli activation (Caspary et al., 2007; Jacoby et al., 2009; Lai et al., 2011). Changes in lipid metabolism are also likely involved as ARL13B directly binds both INPP5E (Qiu et al., 2021), a phosphoinositide 5’-phosphatase, and ARL3, to activate it and thereby promote release of INPP5E from its transporter PDE6D.

Coordinated protein traffic to and within the cilium is essential for both its formation (ciliogenesis) and function (ciliary signaling). Small proteins (<∼100kDa) can diffuse across the barrier at the base of the cilium, the transition zone (Dishinger et al., 2010; Nachury and Mick, 2019). The passage of other proteins through the transition zone is tightly controlled by the IFT protein complexes, IFT-A and IFT-B, and the BBSome (Lechtreck, 2015; Wingfield et al., 2018). Cilium assembly and length are regulated in large part by intraflagellar transport (IFT) machinery, with IFT-A and IFT-B complexes involved in retrograde and anterograde traffic, respectively (Nachury, 2018; Wang et al., 2021). These multi-subunit complexes are required for the regulated entry and export of proteins from cilia, acting in concert with another protein complex, the BBSome, at the base of cilia in the transition zone. IFT-A is also required for efficient traffic of ciliary membrane cargos to the cilium, acting as a vesicle coat complex between the Golgi and cilia (Quidwai et al., 2021). While the core and peripheral protein components of IFT-A and IFT-B have been established (e.g., IFT-A is comprised of Ciliary protein content is further regulated by targeting of newly synthesized proteins from the ER to the Golgi to the cilium and recycling of membrane proteins from the plasma membrane to the ciliary membrane either through lateral diffusion or recycling through endosomes, though these pathways are less well understood than traffic within the cilium itself (Carter and Blacque, 2019; Nachury, 2018).

To begin testing the hypothesis that ARL16 is important to ciliary biology and to search for additional roles in cell biology we knocked out the gene in immortalized mouse embryonic fibroblasts (MEFs). These lines display defects in ciliogenesis, ciliary protein traffic, and ciliary signalling, confirming the phylogenetic prediction and experimentally linking a fourth ARL to cilia. Furthermore, the ciliary traffic defects observed in *Arl16* KO cells appear to be linked to, and possibly the direct result of, changes in traffic of key ciliary proteins through the Golgi. Thus, these data provide an initial characterization of the cellular functions of ARL16 linked to cilia, as well as a role in protein export from the Golgi.

## RESULTS

### ARL16 is a divergent member of the ARF family

ARL16 is among the least characterized GTPases in the ARF family despite its predicted presence in the last eukaryotic common ancestor (Vargova et al., 2021). Sequence alignments reveal ARL16 to be among the most divergent members of the family in mammals, sharing only ∼27% identity to ARF1 (compared to >60% identity among the ARFs and >40% identity among most ARLs) (Fig. S1). Mammalian tissues widely express ARL16 mRNAs, based on searches of RNA and proteome databases, including NCBI Gene and GTexPortal. ARL16’s length is unusually variable amongst mammals, with two variants expressed in humans (Fig. S1 A). The shorter human transcript (NM_001040025.3), encoding a 173 residue protein, is common among mammals, though a longer human isoform of 197 residues is also reported (NM_001040025.2). The longer variant corresponds to an mRNA starting from an upstream transcription initiation site, extending the 5’ end to include an alternative upstream initiation codon. In mice, a frame-shifting indel precludes a similar extension of the coding sequence of the ARL16 gene (not shown), thus mice express only the 173 residue protein. The prevalent, shorter form results in a truncated N-terminus, compared to every other ARF family member, with the G-1 motif (GXXXGKT) beginning at residue 6, in contrast to residue 24 in ARF1 (Fig. S1 A). We predict this difference in the N-terminus has potentially important functional consequences for two reasons. First, the N-termini of ARF family GTPases make direct contacts with binding partners/effectors, effectively serving as a conformationally sensitive “switch III” (Zhang et al., 2009). Also, the shorter form has a cysteine at residue 2 that is predicted to be S-palmitoylated (e.g., by the online prediction tool CSS-Palm (http://csspalm.biocuckoo.org/online.php)). In contrast, the N-terminal extension in the longer isoform moves the cysteine well into the protein where it is no longer predicted to be palmitoylated.

ARL16 proteins retain most of the “G-motifs” (highlighted in red in Fig. S1 A and boxed in red in Fig. S1 B) present in ARF and RAS superfamily members. Mammalian ARL16 retained the conserved PTXG (G-2) motif that is critical in interconversion between the active and inactive conformations as well as the consensus G-4 motif (NKXD) found in almost all nucleotide binding proteins. In contrast, the ARL16 G-3 motif is altered from the highly conserved WDXGGQ (in which the glutamine participates directly in GTP hydrolysis) to RELGGC in ARL16 from multiple species, including humans and mice (Fig. S1 A and B). This change suggests the use of an alternative mechanism of GTP hydrolysis and presents a challenge to researchers as it lacks the glutamine (Q71 in ARFs or Q61 in RAS) in this motif that is commonly mutated to generate a “dominant activating” mutant, used to identify novel functions when expressed in cells.

### The phylogenetic profile of ARL16 predicts a cilium-associated function

Using the database of ARF family GTPase sequences from 114 eukaryotic species described previously (Vargova et al., 2021), we examined the distribution pattern of GTPases missing from eukaryotes that lack ciliated (flagellated) cells or stages. This revealed a strong correlation between ciliated species and four ARF family members, ARL3, ARL6, ARL13, and ARL16 (Fig. 1). To further assess this emerging pattern, we exploited existing genome and transcriptome assemblies to check for the presence of ARL16 orthologs in 25 additional eukaryotes (Table S1, Table S2). The expansion of the sampling was guided by an attempt to cover representatives of additional major phylogenetic lineages of eukaryotes missing in the previous study and to include additional instances of independently evolved non-ciliated taxa. This expanded analysis further supported the view of ARL16 as a broadly conserved and ancient ARF family paralog, as virtually all major eukaryote lineages include at least some ARL16-carrying representative (Fig. 1; Table S1). Multiple alignment of the diverse eukaryote ARL16 sequences revealed that the N-terminal truncation characteristic of the shorter human variant (see above) is in fact a rule rather than exception among ARL16 orthologs (Fig. S1 B). The majority of ARL16 proteins across the eukaryote phylogeny are predicted to be S-palmitoylated near the N-terminus (Vargova et al., 2021), but many lack this modification as they do not have any cysteine residues in that region (Fig. S1 B). Interestingly, the G-4 motif, critical for the specificity of binding of guanine nucleotides (Wittinghofer and Vetter, 2011), is abrogated in ARL16 proteins from several unrelated taxa (Fig. S1), suggesting that these proteins may function as ATPases (Leipe et al., 2002). Overall, ARL16 appears to be a rapidly evolving ARF family subgroup with differences in its primary structure suggesting substantial variation in the molecular details of ARL16 functioning among taxa.

**Figure 1.**
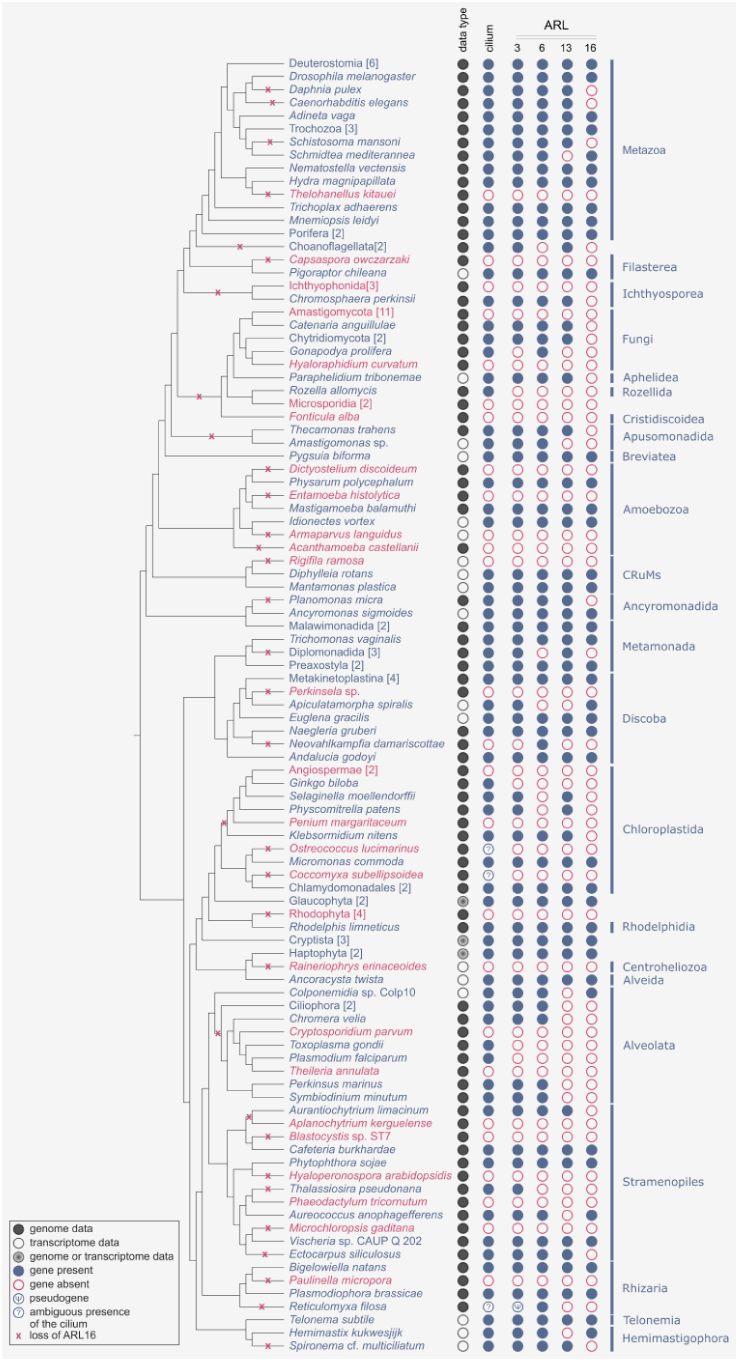
Phylogenetic distribution of ARL16 and the three previously known cilium-associated ARF family members (ARL3, ARL6, and ARL13) and cilia in eukaryotes. The distribution of the four ARF-like genes and cilium (columns) is depicted by filled blue circle (gene/cilium is present) or empty red circle (gene/cilium is absent or not detected). In three species the status of the cilium is ambiguous (see Materials and methods and Table S1). The sequence data type analyzed to establish the presence/absence of the ARLs in the different taxa is indicated (“genome or transcriptome” means that these two types of resources had to be combined to obtain a full set of the orthologs sought). The absence of any of the four ARLs in taxa represented only by transcriptome assemblies must be considered tentative, as the respective genes may be present but not expressed in the stage used for generating the transcriptome data. The schematic phylogeny on the right reflects the current state of knowledge based on multiple phylogenomic analyses. Monophyletic groups of two or more species uniform as to the presence/absence of the cilium and the four focal ARLs are shown as single branches, with the name of the respective broader taxon and the number of taxa included indicated in square brackets (see Table S1 for the full species list). The relationship at the level of the deepest branches, still not completely settled, is adopted from the most recent and comprehensive phylogenomic analysis (Tice et al., 2021), whereas the root of the tree is indicated following the rooting hypothesis by Derelle, et al. (Derelle et al., 2015). Independent losses of the ARL16 gene as inferred by parsimony reasoning from its distribution of extant taxa are mapped onto the phylogenetic tree.

In contrast to some other ancestral ARF family paralogs, ARL16 does not appear to have undergone lineage-specific gene duplications as there are few paralogs with (potentially) differentiated function. The only two taxa with more than one ARL16 gene (the rotifer *Adineta vaga* and the parabasalid *Trichomonas vaginalis*, each have two paralogs) both accumulate duplicated genes due to whole-genome or massive segmental duplications, respectively (Barratt et al., 2016; Flot et al., 2013). Thus, the most conspicuous aspect of ARL16 evolution in eukaryotes is frequent independent loss. The distribution of ARL16 orthologs across the eukaryote tree implies >30 independent loss events, scattered across different time points (affecting recently evolved taxa as well as ancient radiation of major eukaryote taxa; Fig. 1). Notably, the expanded sampling confirmed the initial observation that ARL16 occurs only in eukaryotes for which the ability to form cilia is directly documented or at least predicted by genomic evidence (*i*.*e*., presence of genes encoding typical ciliary proteins; Fig. 1, Table S1). The loss of ARL16 in relation to the loss of the cilium is non-random with high statistical significance (best tail p > 4⨯ 10^−6^, pairwise comparisons as implemented in Mesquite package, see Materials and Methods). However, ARL16 is also absent from various taxa that retained a cilium (Fig. 1; Table S1). Based on these observations, we hypothesize that the cellular role of ARL16 is related to the cilium, more specifically to be dependent on it but not necessarily essential for the function of the cilium as such. To provide a context for the putative association of ARL16 and cilia, we investigated the distribution of ARL3, ARL6, and ARL13 orthologs in the same taxa as we did for ARL16. Like ARL16, orthologs of ARL3, ARL6, and ARL13 are virtually always absent from taxa that do not form cilia (the sole exception being two highly divergent putative ARL6 orthologs in the heterolobosean *Neovahlkampfia damariscottae* that may have been recruited for a novel cilium-independent function). Thus, ARL3, ARL6, and ARL13 can be missing from ciliated taxa but this occurs less frequently than for ARL16 (Fig. 1, Table S1). Altogether, these comparative genomic analyses provide a strong case for the hypothesis that ARL16 is functionally linked to the cilium.

### ARL16 localizes to primary cilia in cultured human RPE cells and photoreceptor cells of the human retina

ARL proteins frequently are found in multiple locations in cells reflecting that a single ARL can perform multiple functions (Sztul et al., 2019). ARL16 was previously found in cytosol in HEK293 and HeLa cells (Yang et al., 2011). We performed immunofluorescence studies of endogenous ARL16 in MEFs and RPE1 cells. Despite the fact that the human and mouse ARL16 proteins (173 residue variant) share 86% identity, the only commercially available ARL16 antibody is specific to the human protein. Thus, we could not identify specific staining of ARL16 in MEFs or in cryosections through murine retinae (data not shown). In contrast, staining of human retinal pigmented epithelial (hTert-RPE1 (RPE1)) cells revealed ARL16 localizes along the ciliary axoneme in a punctate manner (Fig. 2 A). ARL16 also localized to both the cytosol and mitochondria, as evidenced by its diffuse staining across the cell and co-localization with the mitochondrial protein HSP60, respectively (Fig. S2 A).

**Figure 2.**
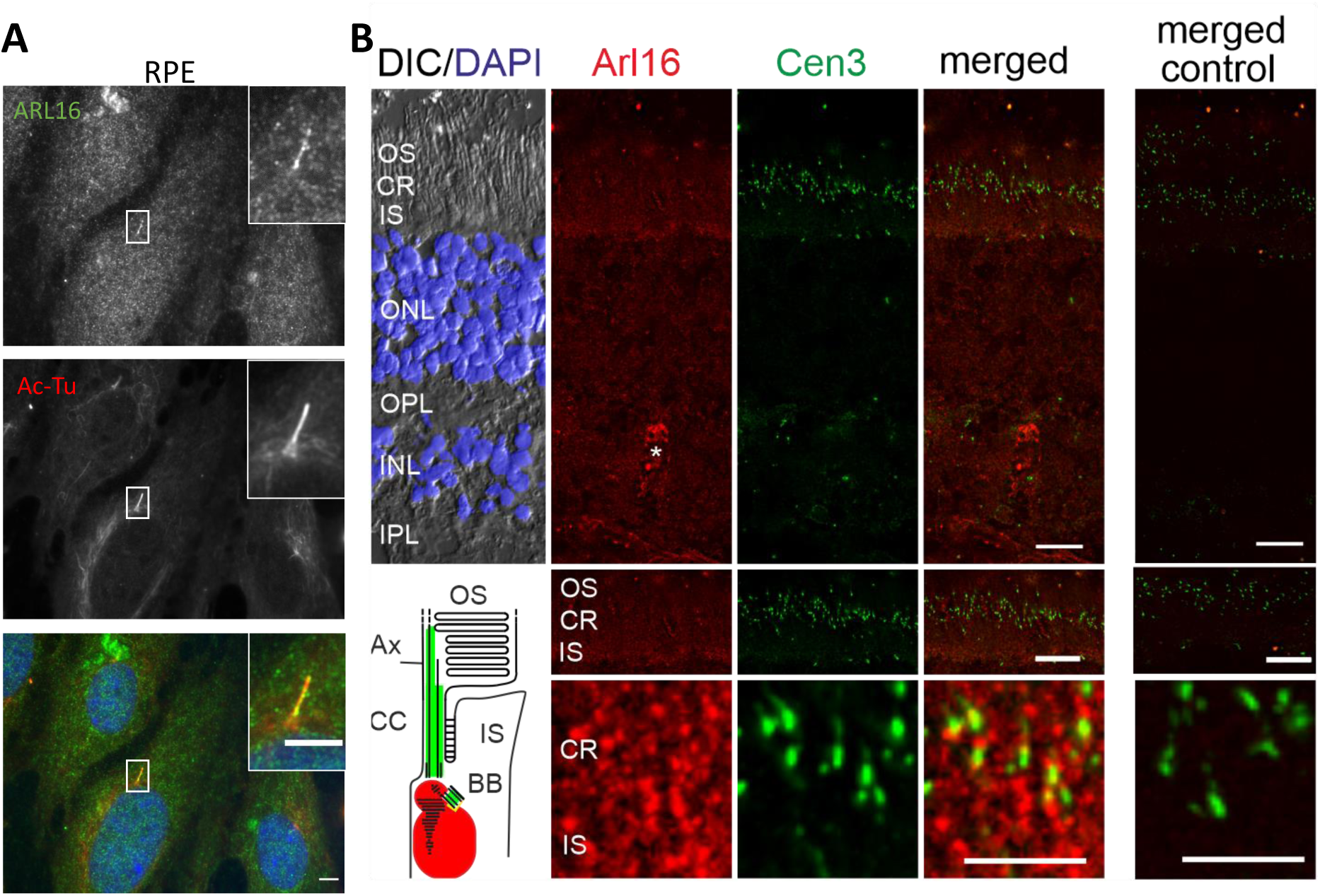
Characterization of endogenous ARL16 localization in RPE1 cells and human retina. **(A)** ARL16 localizes to cilia in RPE1 cells, as observed by immunocytochemistry after PFA fixation (see *Materials and Methods)*. RPE1 cells were serum starved for 48 hrs and stained for ARL16 (green), Ac-Tu (red), and Hoechst (blue). **(B)** Indirect co-immunolabeling of ARL16 (red) and centrin 3 (green), a common marker for the connecting cilium (CC) and the basal body (BB), of a human retina revealed immunofluorescence of ARL16 in the ciliary region (CR) and the inner segment (IS) of retinal photoreceptor cells. Photoreceptor outer segments (OS), DAPI-stained outer and inner nuclear layer (blue, ONL, INL), and outer and inner plexiform layer (OPL, IPL) did not show substantial staining. Asterisk (*) indicates anti-ARL16 blood vessel staining. No staining was observed in controls lacking ARL16 antibodies. Scheme of ARL16 localization in photoreceptor cells. ARL16 is localized to periciliary/basal body region (BB) and in the IS of human photoreceptor cells. Scale bars: A, 10 µm. B, 15 µm and 5 µm (higher magnification).

To further examine the localization of ARL16 in MEFs and RPE1 cells, we generated plasmids that direct human ARL16 or mouse ARL16-myc expression. We generated homologous plasmids for the long or short form of human ARL16. Upon expression of the 173 residue human or mouse proteins in MEFs, we observed diffuse ARL16 staining in the cytosol (Fig. S2 B). In contrast, the 197 residue human variant displayed little evidence of diffuse, cytosolic localization. Instead, tubular and punctate staining co-localized strongly with HSP60 at mitochondria, consistent with endogenous staining of RPE1 cells after methanol fixation (Fig. S2 A). Thus, different length human variants localize quite distinctly when exogenously expressed. Because the endogenous staining of ARL16 in RPE1 cells displays prominent mitochondrial localization, we believe that these cells express at least some of the longer form of ARL16. Despite the presence of endogenous ARL16 in cilia in RPE cells, we did not find evidence of the exogenous protein localizing to cilia in MEFs.

We also analyzed ARL16 localization in photoreceptor cells in the retina, a commonly studied ciliary model tissue. The light sensitive outer segment resembles a highly modified but well characterized primary cilium (May-Simera et al., 2017; Roepman and Wolfrum, 2007). We used indirect immunofluorescence staining on a human donor retina as previously reported (Davidson et al., 2013; Turn et al., 2021). Figure 2 B shows double labeling of ARL16 and centrin 3, a marker for the connecting cilium as well as the mother (basal body) and daughter centrioles of photoreceptor cells in human retinae. Prominent ARL16 immunofluorescence labelled the ciliary region (CR) and the inner segment of the photoreceptor layer. Other retinal layers, such as the outer and inner nuclear layer (ONL, INL), outer and inner plexiform layer (OPL, IPL) where retinal synapses are localized, and the ganglion cell layer (GCL) showed no substantial staining. Higher magnification of the ciliary region revealed the localization of ARL16 in the periciliary region at the base of the connecting cilium (stained by anti-centrin 3) in human photoreceptor cells (Fig. 2 B, lower panels). The centrin-3 co-staining further showed that ARL16 labeled the basal body next to the daughter centriole. The inner segment where all biosynthetic active organelles such as the Golgi and ER are found, also showed prominent ARL16 immunofluorescence. Similar localizations were previously found for ciliary molecules which participate in Golgi functions and/or Golgi and molecular transfer processes to ciliary base (Sedmak and Wolfrum, 2010). The localization at the base of the connecting cilium is in line with other GTPases of the ARF superfamily like ARL2 and the ARF GAP ELMOD2, which also localize to ciliary rootlets in the inner segment (Turn et al., 2021).

### CRISPR/Cas9 Knockout of *Arl16* in MEFs

To test the prediction of a role for ARL16 in ciliary function, and to assess broader questions of its function in cells, we used CRISPR/Cas9 to introduce frame-shifting mutations into the open reading frame of *Arl16* in MEFs. These cells are frequently used in cell biological studies, easily imaged due to their flat morphology, commonly used in cilia research, and diploid (which facilitates genome editing analyses). We used two guides to target Cas9 to exon 2, just downstream of the ATG at the end of exon 1 in *Arl16* and screened clonal lines for indels by DNA sequencing of targeted regions. We sought frameshifting mutations predicted to cause premature translation termination and loss of protein function. The lack of a mouse antibody precluded us from directly testing for absence of protein expression. Although there is strong evidence (Smits et al., 2019) that cells may express mutated proteins resulting from use of downstream start sites or alternative splicing, the small size of the *Arl16* open reading frame and presence of multiple, essential G-motifs along the length of the protein makes expression of a functional protein after introduction of indels near the N-terminus highly unlikely. Thus, we refer to these as knockout (KO) lines. We generated five *Arl16* KO clonal lines from two different guides that we used in these studies (alleles are shown in Fig. S3 A). Because we only found minor differences between the KO lines, data are presented in aggregate. As a further test of specificity and to ensure against off-target effects influencing our phenotyping, we performed rescue experiments in which we exogenously expressed ARL16-myc in KO and WT lines and assessed the reversal of phenotypes arising from the KOs.

Because almost nothing is known about ARL16 functions, and to obtain an unbiased overview of the effect of ARL16 loss on cells, we screened markers of major organelles and the cytoskeleton to look for differences between the *Arl16* KO lines and the parental WT MEFs. No obvious differences were observed in mitochondrial morphology or distribution (HSP60), f-actin (FITC-phalloidin), nuclear number, size, or morphology (Hoechst), or microtubule networks (α/β-tubulin) when cells were fixed and stained as indicated (Fig. S3 B). However, these screens revealed a number of cellular defects in *Arl16* KO lines, which we describe below.

### *Arl16* KO cells have reduced ciliation and increased ciliary length

In light of the phylogenetic prediction of a role for ARL16 at cilia and its localization there, we examined *Arl16* KO cilia. We used acetylated tubulin (Ac-Tu) staining throughout to mark the ciliary axoneme after inducing ciliation in cultured cells. All five of the *Arl16* null lines displayed reductions in the percentage of ciliated cells, compared to WT MEFs (Fig. 3 A, B). When the data from the five KO lines were pooled, the KO’s had about 50% as many ciliated cells as WT after 24 hrs. However, at later time points the differences between WT and KO lines decreased, down to ∼25% lower at 72 hrs of serum starvation and without reaching statistical significance. Thus, there is a decrease in ciliation compared to WT.

**Figure 3:**
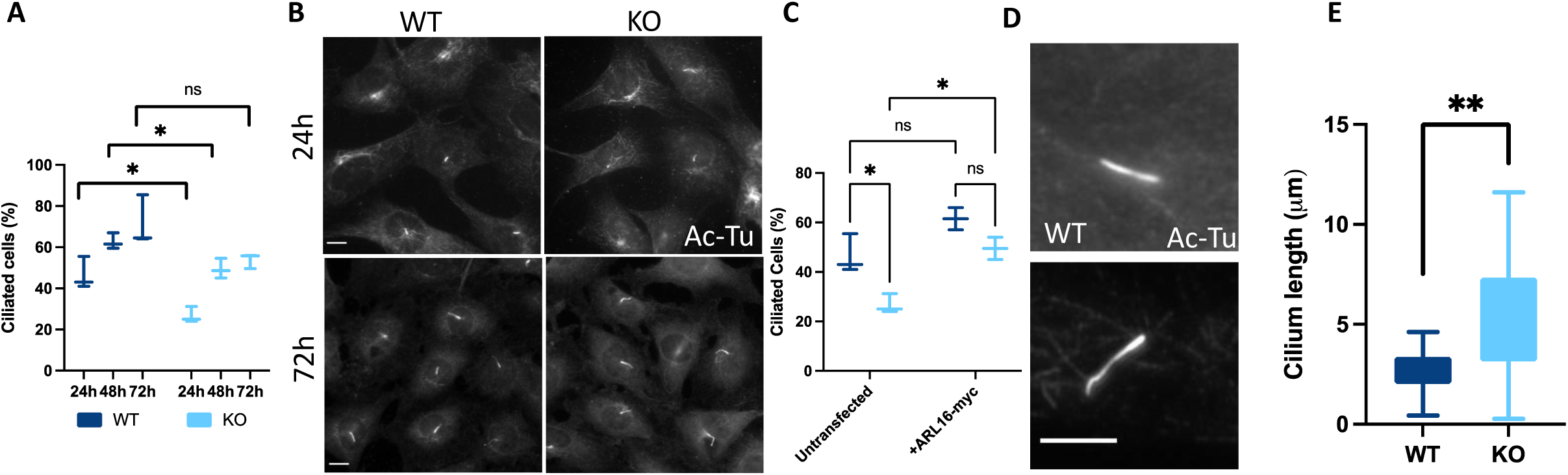
Deletion of *Arl16* causes defects in ciliogenesis. (**A**) WT and *Arl16* KO MEFs were serum starved with percentage of ciliated cells manually counted. For each cell line (n=2 WT and n=5 KO), 100 cells were scored at each time point in triplicate and averaged. Multiple unpaired t-tests, false discovery rate Q= 1%, Two-stage step-up (Benjamini, Krieger, and Yekutieli). 24h p=0.017503, 48h p=0.020865, 72h p=0.076598 (**B**) *Arl16* KO cells fail to ciliate as efficiently as WT cells, particularly at earlier timepoints after serum starvation. WT and *Arl16* KO MEFs serum starved for 24h or 72h and stained for Ac-Tu. Scale bar = 10 μm, 60x. (**C**) Expression of ARL16-myc restores ciliation in *Arl16* KO cells after 24h of serum starvation. Values for untransfected cells are equivalent to the data from panel A at 24h of serum starvation. Ciliation of transfected cells was scored as in panel A (N=2×50 transfected cells from one WT line and 2 KO lines). Two way ANOVA Tukey’s multiple comparisons (Untransfected WT vs KO p=0.0315, WT transfected vs untransfected p=0.1349, KO Untransfected vs Transfected p=0.0277, Transfected WT vs KO p=0.3133) (**D**) *Arl16* KO cilia are longer than WT cilia. Cells were serum starved for 48h before fixing and staining for Ac-Tu, and imaged at 100X. Scale bar 5 µm. (**E**) Cilium lengths were measured in WT and *Arl16* KO cells using the CiliaQ Plugin in FIJI. N=26 cilia for 1 WT and 2 KO lines. T-test P=0.0023.

To ensure that the changes in ciliation were due to the loss of ARL16 and not off-target effects, we expressed ARL16-myc in WT and *Arl16* KO cells. One day after transfection, we serum starved for 24 hrs and scored the percentage ciliation in cells expressing ARL16-myc. Interestingly, the percentage of WT cells with cilia increased almost 20% upon ARL16-myc expression, though this difference did not reach statistical significance (Fig. 3 C; p>0.05). Expression of ARL16-myc in *Arl16* KO lines (n=2 lines) resulted in a larger increase in ciliation, to levels approaching, and not statistically different from, those seen in WT cells also expressing ARL16-myc. Thus, expression of ARL16-myc reversed the decrease in ciliation observed in *Arl16* KO lines, supporting a role for ARL16 in ciliogenesis.

It was also evident that cilia in *Arl16* KO cells are longer than those in WT cells. Using the CiliaQ Plugin in FIJI (Hansen et al., 2021), we measured cilia lengths after 48 hrs of serum starvation (Fig. 3 D, E). On average, cilia in *Arl16* KO lines (5.09 µm) are ∼90% longer than in WT (2.67 µm) cells, based upon measurement of Ac-Tu staining.

To gain insights into likely causes of the deficits in ciliation in *Arl16* KOs, we screened markers of ciliogenesis for differences compared to WT MEFs. CEP164 recruitment to mother centrioles is an early step in ciliogenesis and required for cilium formation, while CP110 uncapping is a later step just prior to axoneme elongation (Cajanek and Nigg, 2014; Yadav et al., 2016). We examined these proteins in WT and *Arl16* KO MEFs and found no statistically significant differences in either CEP164 recruitment or CP110 uncapping after 24 hr of serum starvation (Fig. S4 A, B). Thus, we conclude that the defect in ciliogenesis observed in the *Arl16* KO MEFs is downstream of CP110 uncapping.

Ciliary rootlets surround the basal body, are important in centrosome cohesion, and often project a foot or extension that helps stabilize cilia (Hossain et al., 2020; Yang et al., 2005; Yang and Li, 2006; Yang et al., 2002). Thus, we stained WT and *Arl16* KO cells for rootletin, the primary component of rootlets. The absence of ARL16 strongly correlated with rootlet fragmentation (Fig. S4 C, D). However, in contrast to results observed in *Elmod2* KOs, we did not observe an increase in centrosome separation in *Arl16* KOs, based upon staining for γ-tubulin to mark centrosomes (Fig. S4 E).

### *Arl16* KO leads to loss of several proteins from cilia

While cilia could be readily identified by Ac-Tu staining in all cells, ciliary ARL13B staining in *Arl16* KOs was dramatically reduced compared to that seen in WT cells (Fig. 4 A, B). Because recruitment of ARL13B to cilia is required for ciliary accumulation of the other ciliary membrane proteins, specifically ARL3 and INPP5E (Gigante et al., 2020), we assessed the endogenous levels and localizations of each of these proteins to the cilia of *Arl16* KO cells. We readily detected both ARL3 and INPP5E in cilia of control MEFs, but neither ARL3 nor INPP5E were found in *Arl16* KO cilia (Fig. 4 C-F). Finally, we assessed the localization of the ciliary transmembrane protein adenylyl cyclase 3 (AC3) and found that AC3 is also lost from cilia of *Arl16* KOs.

**Figure 4:**
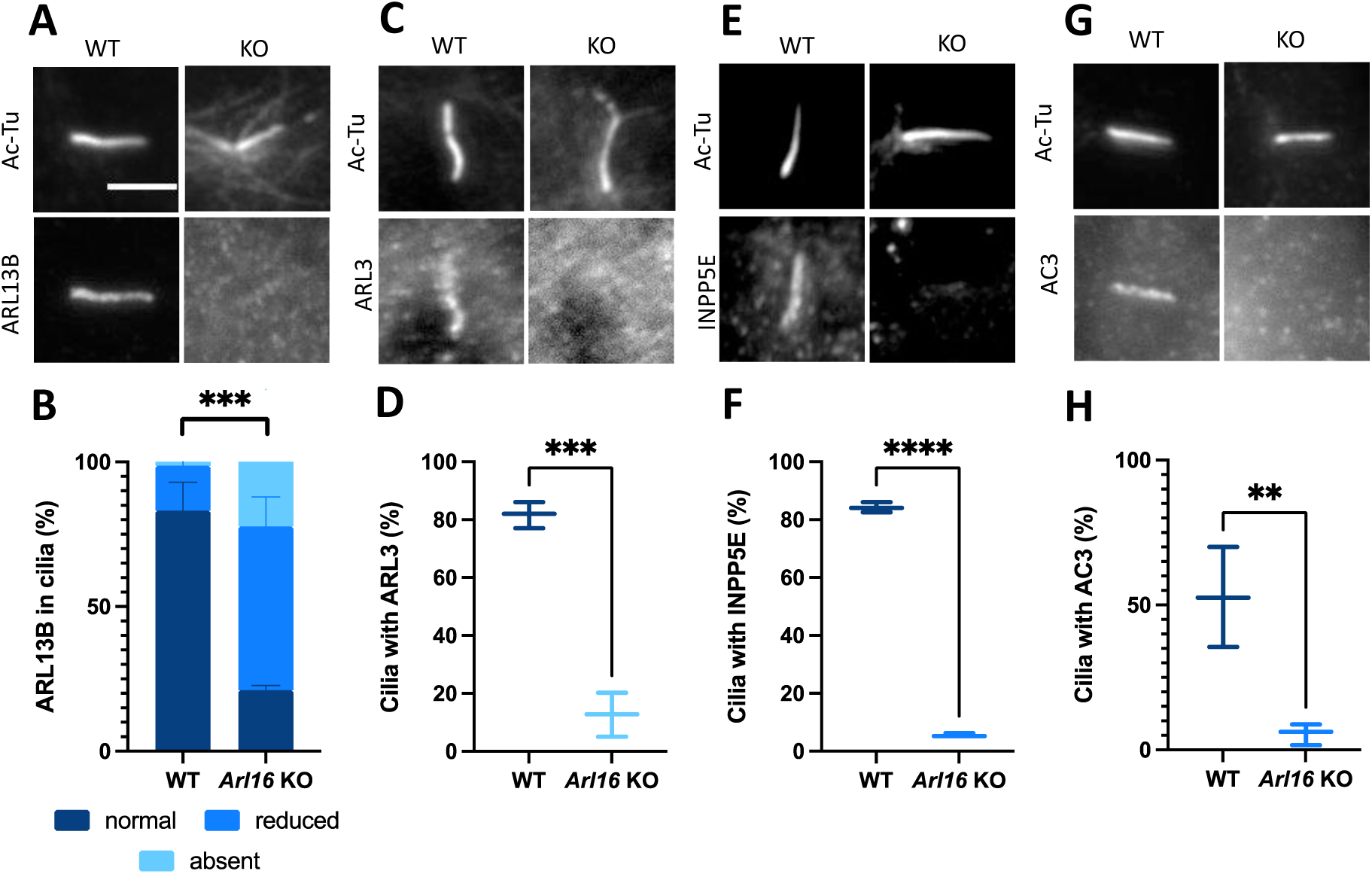
*Arl16* KO cells have reduced recruitment of several ciliary proteins. **(A)** ARL13B levels are reduced in *Arl16* KO cilia. Cells were serum starved for 48h and stained for Ac-Tu and ARL13B. **(B)** Quantification of ARL13B positive cilia in WT and *Arl16* KO lines. N=3×100 cilia X 5 KO lines or 2 WT lines. Cilia were identified using the Ac-Tu channel. Presence of ARL13B was scored by eye. Normal ARL13B staining was defined as staining that was readily apparent independent of the Ac-Tu channel. Reduced staining was only apparent after identification of the cilium using the Ac-Tu channel. T-test. P=0.0002. **(C)** ARL3 is reduced in *Arl16* KO cilia. Cells were serum starved for 48h and stained for Ac-Tu and ARL3. **(D)** Quantification of C. N=3×100 cilia X 5 KO lines or 2 WT lines. Cilia were identified using the Ac-Tu channel. Presence of ARL3 was scored by eye. T-test p=0.0002 **(E)** INPP5E is absent from *Arl16* KO cilia. Cells were serum starved for 48h and stained for Ac-Tu and INPP5E. **(F)** Quantification of E. N=3×100 cilia X 5 KO lines or 2 WT lines. Cilia were identified using the Ac-Tu channel. Presence of INPP5E was scored by eye. T-test p<0.0001. **(G)** AC3 is reduced in *Arl16* KO cilia. Cells were serum starved for 48h and stained for Ac-Tu and AC3. **(H)** Quantification of G. N=3×100 cilia X 5 KO lines or 2 WT lines. Cilia were identified using the Ac-Tu channel. Presence of AC3 was scored by eye. T-test p=0.0098. Scale bar=5 µm 100x for all images.

To determine if there was a generalized disruption in ciliary protein traffic, or a more specific loss related to ARL13B and ARL3, we expressed GFP-tagged somatostatin receptor (SSTR3-GFP) and assessed its ciliary localization in WT and *Arl16* KO MEFs. We observed similar SSTR3-GFP signal in the cilia of both *Arl16* KOs and WT controls (Fig. S4 F). This suggests that import of certain proteins is unaltered in cells lacking ARL16, though results from the exogenous expression of a tagged protein carries with it known caveats. We also examined a protein marker of the transition zone, CEP290. In both WT and *Arl6* KO cells, CEP290 localizes to the base of the cilium, indicating no evident defects in its localization upon ARL16 ablation (Fig. S4 G). Thus, there is selective loss of a subset of ciliary proteins in cells lacking ARL16.

### Shh signaling is defective in *Arl16* KO cells

In light of the loss of ARL13B from cilia in *Arl16* KO lines, and its proven role in ciliary signaling, yet ability to signal from outside cilia, we investigated whether ARL16 regulates Sonic Hedgehog (Shh) signaling (Bay et al., 2018; Caspary et al., 2007; Gigante et al., 2020; Larkins et al., 2011). Treatment of MEFs with Shh causes changes in the ciliary protein content, as well as processing of Gli transcription factors that ultimately result in changes in nuclear transcription, including that of *Ptch1* and *Gli1* mRNAs (Li et al., 2021). Smoothened (SMO) dynamically localizes to the ciliary membrane in response to Shh stimulation, with SMO absent under control conditions and enriched upon stimulation with Shh ligand (Corbit et al., 2005). Cells were treated with Shh conditioned medium with serum starvation for 24 hr and then stained for SMO. SMO robustly increases in WT cilia but fails to do so in *Arl16* KOs (Fig. 5 A, B).

**Figure 5:**
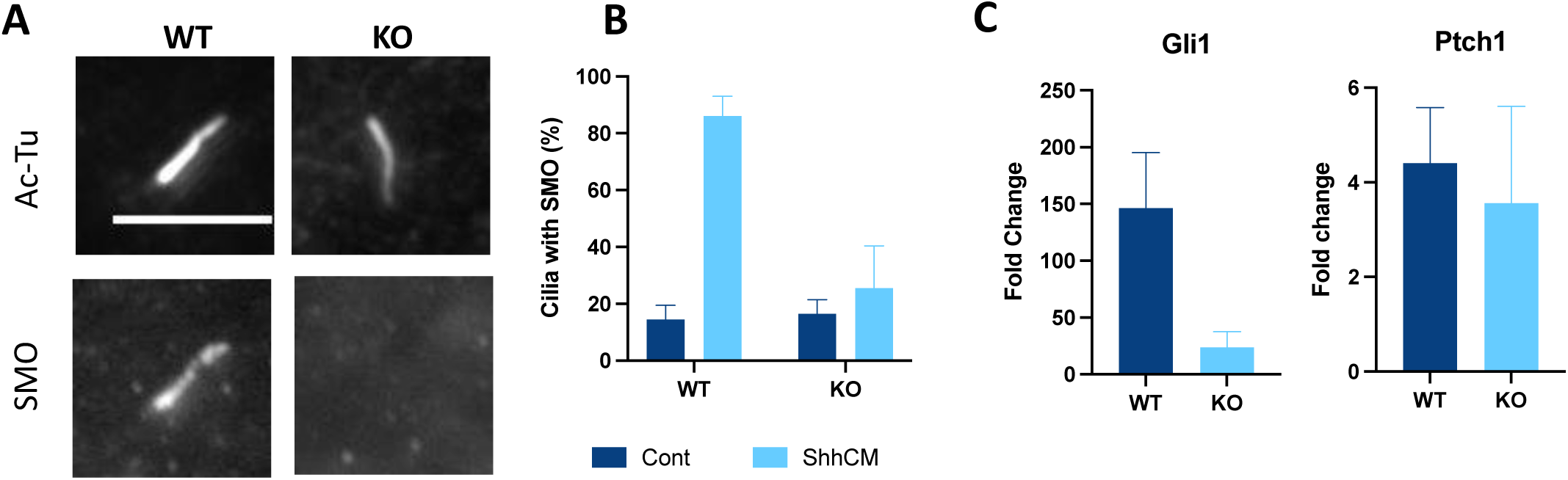
Arl16 KO cells have defects in Hh signaling. (**A**) SMO is lost from cilia in *Arl16* KO lines. Cells were serum starved for 48h and stained for Ac-Tu and SMO. (**B**) Quantification of A. N=2×100 cilia X 2 KO lines and 1 WT line per condition. (**C**) *Arl16* KO MEFs show reduced Shh-stimulated Gli1 and Ptch1 transcriptional response compared to WT cells. Cells were collected 24 h after Shh treatment, and levels of Gli1 and Ptch1 mRNA were determined using qPCR. Bar graphs indicate normalized mRNA expression with data presented as mean fold change ± SEM. N=2 WT lines and 4 KO lines.

To monitor transcriptional changes resulting from treatment of MEFs with Shh ligand, we performed qPCR to measure *Ptch1* and *Gli1* mRNA levels (two well-known targets increased by Shh signaling) after Shh stimulation. We observed increases in the levels of each of these mRNAs in WT cells in response to Shh stimulation. In contrast, the responses were markedly blunted in *Arl16* KO cells (Fig. 5 C). Thus, ARL16 is also required for two well characterized responses to Shh; SMO recruitment and transcriptional changes in target genes.

### IFT140 is absent from cilia of *Arl16* KOs

We also analyzed the localization of several IFT components in *Arl16* KO cells. In WT cells, IFT140 is observed at the ciliary base and along the length of the cilium. In marked contrast, IFT140 associated with the cilium or base was strongly reduced in *Arl16* KO lines (Fig. 6 A). IFT140 is one of three core subunits of the IFT-A complex so its absence suggested the possibility of the absence of this entire complex. Surprisingly, when we examined another core IFT-A component, IFT144, we observed its presence in the cilium with no evidence of changes between WT and *Arl16* KO cells (Fig. 6 B). While optimizing staining protocols for IFT140, we found that IFT140 also localizes to rootlets, displaying strong overlap with rootletin staining in both WT and *Arl16* KO cells (Fig. S4 H), even after fragmentation of rootlets in the KO cells.

**Figure 6:**
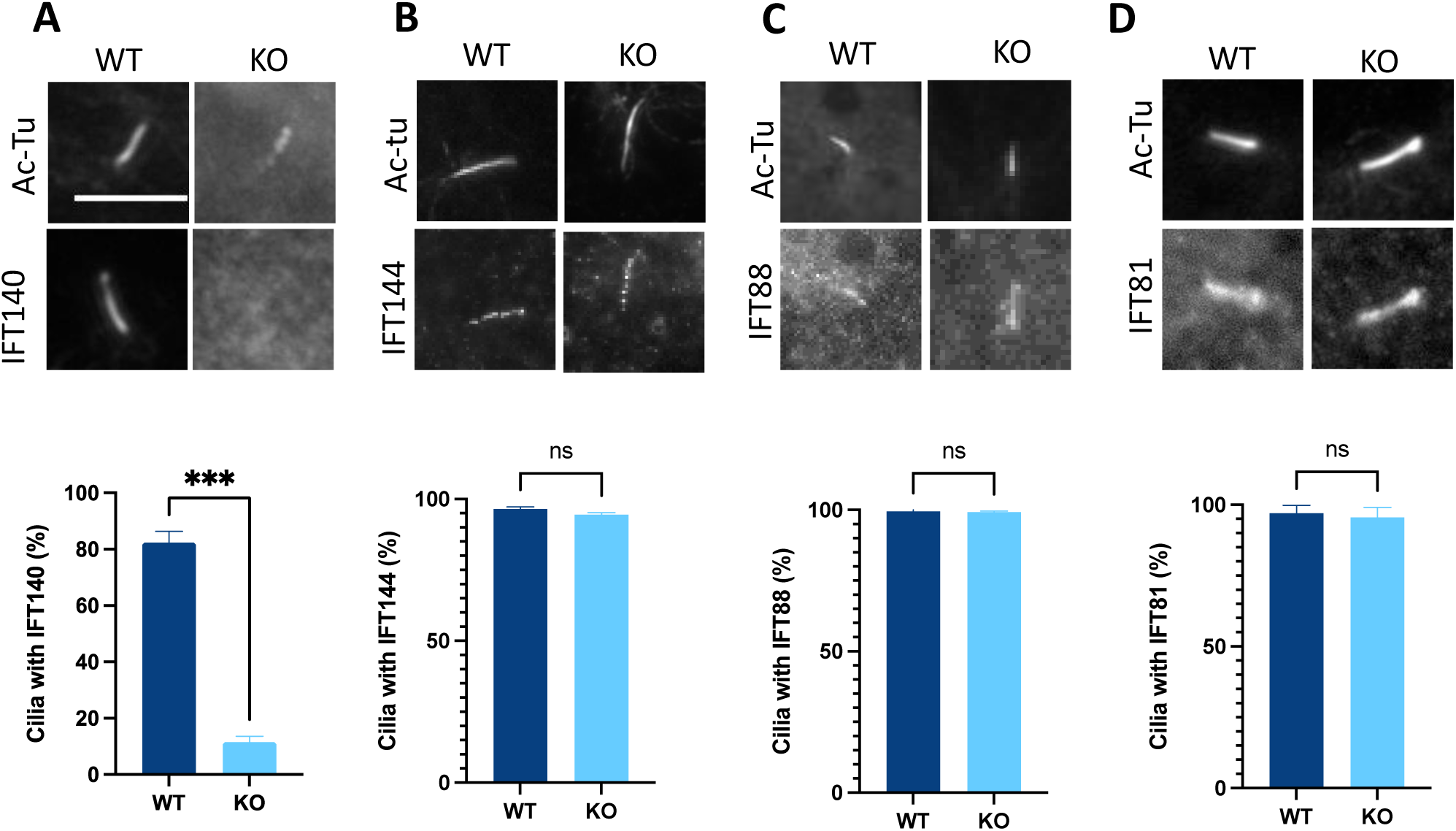
IFT140 is lost from cilia in *Arl16* KO cells, but other IFTs are unchanged. In each case, cells were serum starved for 48 h and then fixed and (top) stained with Ac-Tu and either IFT140 (A), IFT144 (B), IFT88 (C), or IFT81 (D) and imaged at 100x magnification. Quantification was performed in each case in duplicated for 100 cilia in WT (N=2 cell lines) and Arl16 KO lines (N=2 lines). (**A**) IFT140 is lost from cilia in *Arl16* KO lines. (**B**) IFT144 staining in cilia is unchanged between WT and *Arl16* KO lines. p=0.1056. (**C**) IFT88 is not altered in *Arl16* KO cilia from that in WT cells. p=0.6985. (**D**) IFT81 is present in *Arl16* KO cilia as it is in WT cilia. p=0.6855. Scale bar = 5 µm.

We also examined effects of *Arl16* KO on core components of the IFT-B complex, which consists of two core complexes (B1-1 and B1-2) and a peripheral complex (Quidwai et al., 2021; Wang et al., 2021; Yang and Huang, 2019). IFT81 (B1-1) and IFT88 (B1-2) localize along the length of the cilium and are often enriched at the base and tip in MEFs (Quidwai et al., 2021). We observed no differences in the localization of either IFT88 or IFT81 in cilia of *Arl16* KO cells (Fig. 6 C, D), with each of them localizing along the length of the cilium in both WT and KO cells. Therefore, we observed changes in the ciliary localization of a single IFT-A core component, IFT140, without apparent disruption of the other IFT-A or IFT-B components.

### IFT140 and INPP5E accumulate at the Golgi in *Arl16* KO cells

While we did not detect IFT140 in *Arl16* KO cilia, we noted instead that IFT140 accumulated in an intracellular membranous structure, which we identified as the Golgi by co-staining with GM130 (Fig. 7 A). This Golgi staining of IFT140 was not evident in WT cells, and expression of ARL16-myc reversed the increased staining of IFT140 at Golgi (Fig. 7 A). The only other IFT protein previously known to be associated with the Golgi is IFT20 (Crouse et al., 2014; Follit et al., 2006; Keady et al., 2011). Consistent with these earlier results, we found that in WT cells, IFT20 localizes to the Golgi and overlaps strongly with GM130 staining. We stained for IFT20 in *Arl16* KO cells and found no differences in its localization compared to WT cells (Fig. 7 B). We examined the potential presence of other IFT-A (IFT144) or IFT-B (IFT88, IFT81) proteins at Golgi and no increases were evident. We then stained for several other Golgi markers (including β-COP, FAPP2, and GBF1) to assess whether overall Golgi structure may be altered in *Arl16* KO cells. Each of these proteins localize to the Golgi in both WT and *Arl16* KO cells, and we observed no differences in Golgi morphology or intensity of any of these markers, based on visual inspection (Fig. S3 B). Thus, while one component from each of the IFT-A and IFT-B complexes, IFT140 and IFT20, respectively, can be found at the Golgi, only IFT140 is increased in its abundance at the Golgi in *Arl16* KOs, while IFT20 appears to be a resident Golgi protein in both WT and *Arl16* KO MEFs.

**Figure 7:**
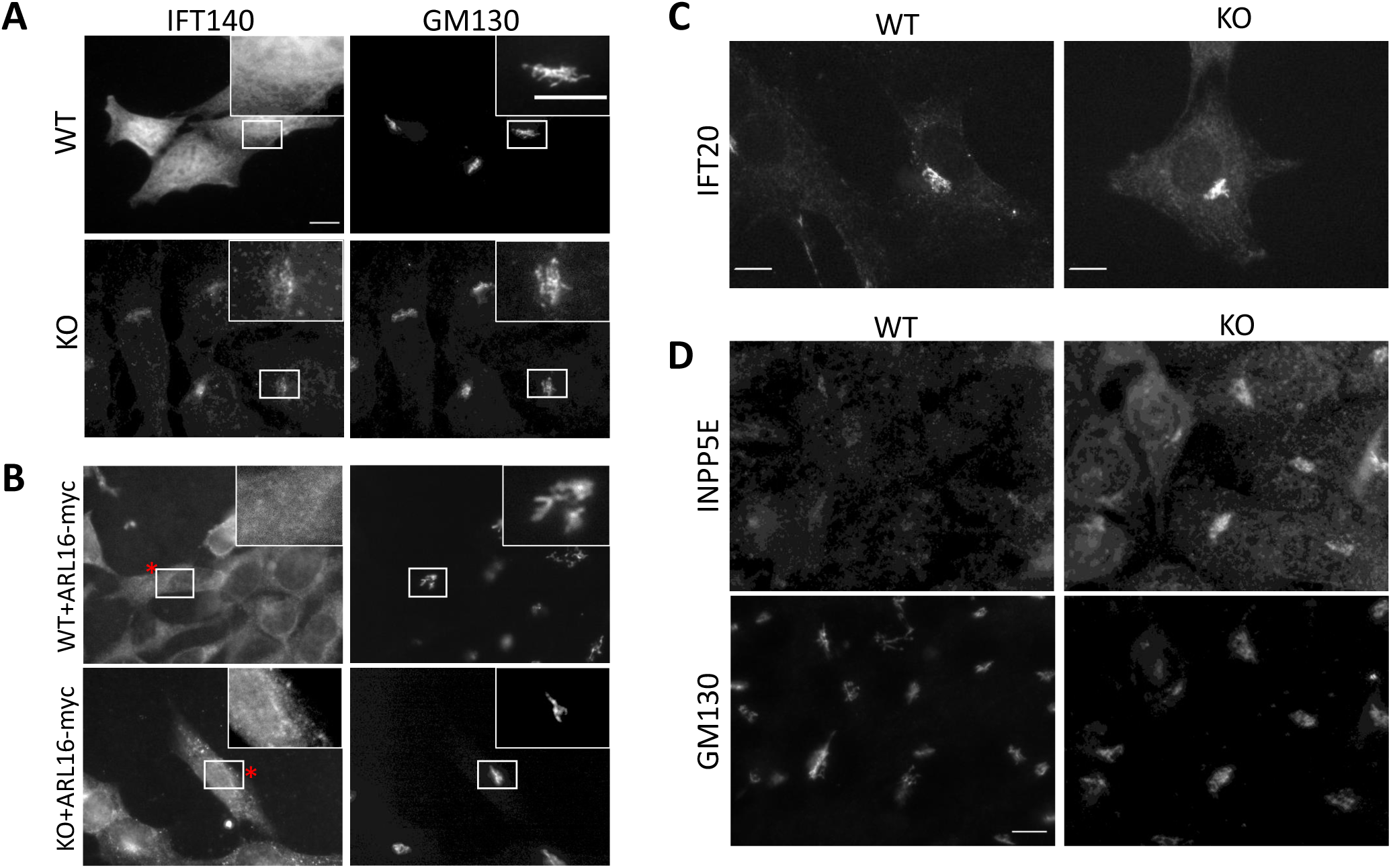
IFT140 and INPP5E accumulate at the Golgi of *Arl16* KO lines. (**A**) Cells were serum starved for 24h and stained for IFT140 and GM130. IFT140 staining is diffuse in WT cells, but clearly enriched at the Golgi in *Arl16* KOs. Asterisks indicate transfected cell. Scale bar 10 µm. (**B**) Expression of ARL16-myc reverses IFT140 accumulation in the Golgi. Cells were transfected with ARL16-myc, serum starved for 24h, and stained for myc, GM130, and IFT140. (**C**) IFT20 localizes to Golgi indistinguishably in WT and *Arl16* KO cells. Cells were serum starved for 24h and stained for IFT20. (**D**) Serum starved cells were fixed and stained after 24 h and stained for INPP5E and GM130. INPP5E co-staining with the Golgi marker GM130 is very strong while there was no evidence of changes in overall Golgi morphology.

Surprised by the accumulation of IFT140 in the Golgi in *Arl16* KOs, we assessed the Golgi localization of the other proteins that were lost from cilia using a variety of staining protocols to see if any others were accumulating there. Staining of INPP5E at cilia is typically performed using PFA fixation and under these conditions we observed it in cilia of WT lines but not in *Arl16* KO lines. However, after methanol fixation, we found INPP5E staining increased at the Golgi in *Arl16* KO but not WT cells (Fig. 7 C). In contrast, neither ARL3 nor ARL13B were observed at the Golgi in either WT or *Arl16* KO lines under any conditions examined.

### INPP5E but not IFT140 is lost from cilia and accumulates in the Golgi of *Pde6d* KOs

The phosphatidylinositol phosphate (PIP) 5’-phosphatase INPP5E is farnesylated at its C-terminus. As a result, its traffic to cilia is thought to be dependent on the prenyl binding protein PDE6D (Cook et al., 2000; Fansa et al., 2016). The prevailing model postulates that PDE6D binds to INPP5E and carries it to the cilium where it is released by ARL3 and retained by ARL13B (Humbert et al., 2012; Kosling et al., 2018; Thomas et al., 2014) A high throughput, yeast two-hybrid screen found that in addition to binding ARL3, PDE6D also binds ARL16 (Rolland et al., 2014). Therefore, we generated *Pde6d* KO MEFs using the same protocols as those to generate *Arl16* KOs (Fig. S5) to compare the phenotypes observed in the absence of *Pde6d* to those observed in the absence of *Arl16*. We generated 5 lines from two guide RNAs in which the *Pde6d* gene contained frameshifting mutations (alleles of lines used in these studies are shown in Fig. S5B).

We then characterized the *Pde6d* KO lines for cilia number and protein composition. We observed no changes in the percentage of ciliated cells or length of cilia between *Pde6d* KO and WT cells (Fig. 7 A,B). We also found no differences in the strength of staining of ARL13B or ARL3 in cilia of *Pde6d* KOs compared to WT cells (Fig. 8 C, D). In contrast, when we examined ciliary staining of INPP5E, it was absent from *Pde6d* and *Arl16* KO lines while present in WT cells (Fig. 8 D). INPP5E was instead increased at the Golgi in *Pde6d* KO cells as in *Arl16* KO cells (Fig. 7 G).

**Figure 8:**
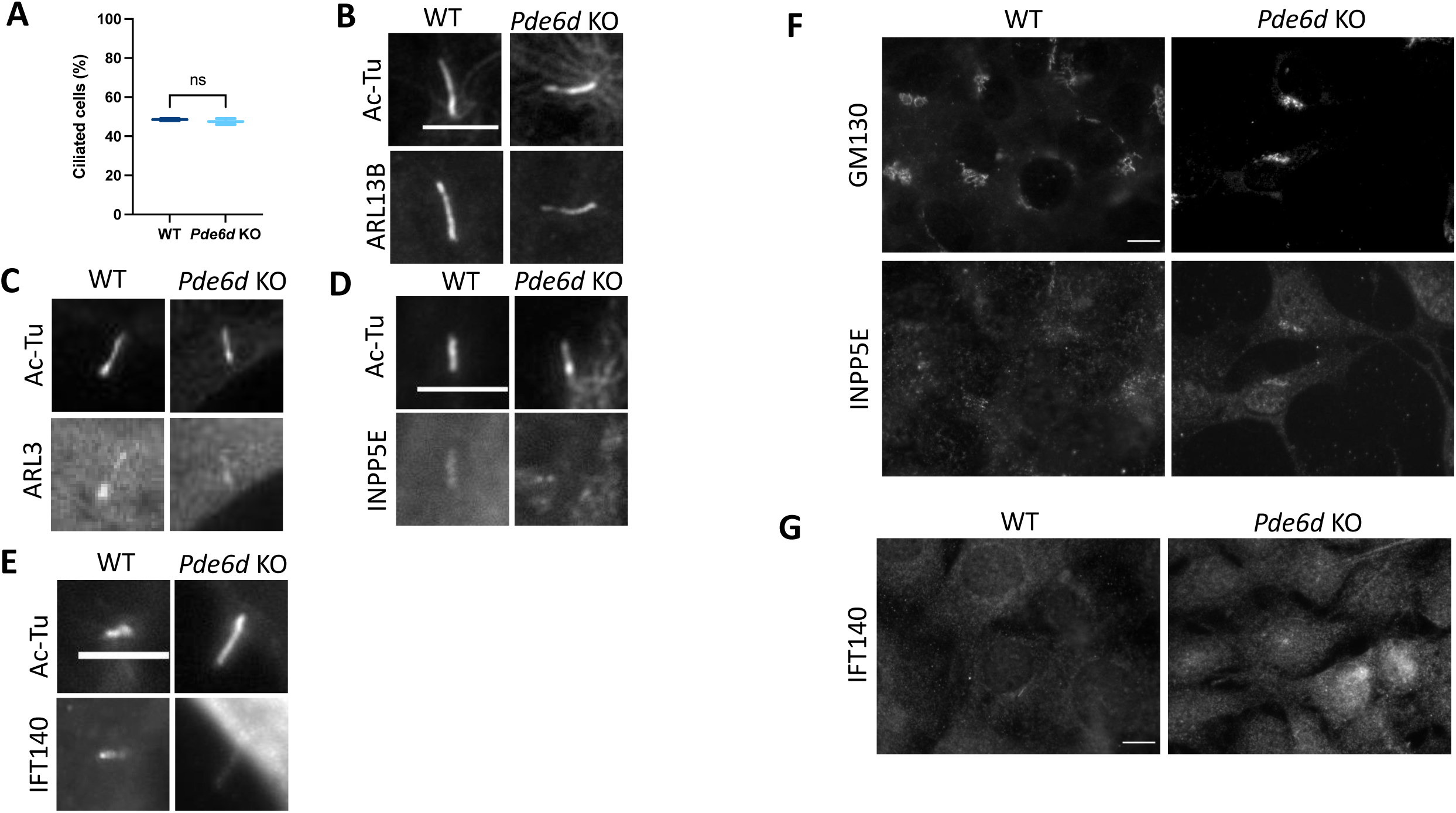
Cells deleted for *Pde6d* display no defects in ciliation but loss of INPP5E from cilia with its accumulation at Golgi. (**A**) Cells were serum starved for 24h, stained for Ac-Tu, and scored for ciliation as described in Materials and Methods. N=2×100 cells for 1 WT and 2 KO lines. T-test p=0.5918. (**B**) ARL13B and (**C**) ARL3 localize normally to *Pde6d* KO cilia. (**D**) INPP5E is lost from *Pde6d* KO cilia. (**E**) IFT140 localizes normally to cilia of *Pde6d* KO cells. (**F**) INPP5E is absent from cilia of *Pde6d* KO lines. Cells were serum starved for 24h and stained for Ac-Tu and INPP5E. (**G**) INPP5E accumulates in the Golgi of *Pde6d* KO lines. Cells were serum starved for 24h and stained for INPP5E and GM130. All images collected using 100x objective and scale bars = 5 µm.

We asked next whether IFT140 is also altered in *Pde6d* KO cells. We found that IFT140 localizes in *Pde6d* KO cilia at comparable levels to WT cilia, and is not enriched at the Golgi (Fig. 8 F, H). Thus, while traffic of both IFT140 and INPP5E to cilia requires ARL16, only traffic of INPP5E appears to be dependent on PDE6D. These results are consistent with previous data showing PDE6D-dependence of INPP5E traffic to cilia, likely the result of the prenylated INPP5E cargo binding the PDE6D shuttle. Further, these data suggest that IFT140 and INPP5E traffic to cilia via different pathways, each of which is dependent upon, or regulated by, ARL16.

## DISCUSSION

We used a combination of phylogenetic analyses, genome editing of MEFs, and cell biological approaches to identify the cellular functions of one of the last uncharacterized members of the mammalian ARF family of regulatory GTPases, ARL16. We identified a strong correlation between the presence of cilia with the presence of ARL16 in genomes across the eukaryotic spectrum, leading us to propose cilia as a major site of action. Consistent with our phylogenetics-based model, we found ARL16 localizes to cilia and basal bodies in RPE and retinal photoreceptor cells, respectively. We generated *Arl16* KO MEF lines to identify gross changes in cellular biology resulting from its loss and to experimentally test its predicted role in one or more aspects of cilia. We found that the loss of ARL16 results in a reduction in ciliation, due to changes in a step downstream of CP110 uncapping. We also observed an increase in average ciliary length and a large decrease in the presence of ARL13B, ARL3, INPP5E, and IFT140 in cilia. However, we believe that the most important function of ARL16 in ciliary biology may lie in its role in the regulation of traffic of ciliary proteins from the Golgi to cilia. We found two such pathways that are compromised when cells lack ARL16, one used by IFT140 and another by INPP5E. Deficiency of either IFT140 or INPP5E at cilia is expected to spawn an unknown number of downstream phenotypes, as they are key regulators of traffic along cilia, of protein export (IFT140) and in the control of ciliary membrane phospholipid composition (INPP5E). We also provide data that support a model for ARL16 acting from multiple sites in cells, including basal bodies and Golgi. Thus, our findings highlight both the importance of ARL16 to multiple processes and sites (summarized in the model shown in Fig. 9) and also the challenges involved in future work aimed at identifying molecular mechanisms for each of its actions, as might be expected for such an ancient and highly conserved GTPase.

**Figure 9:**
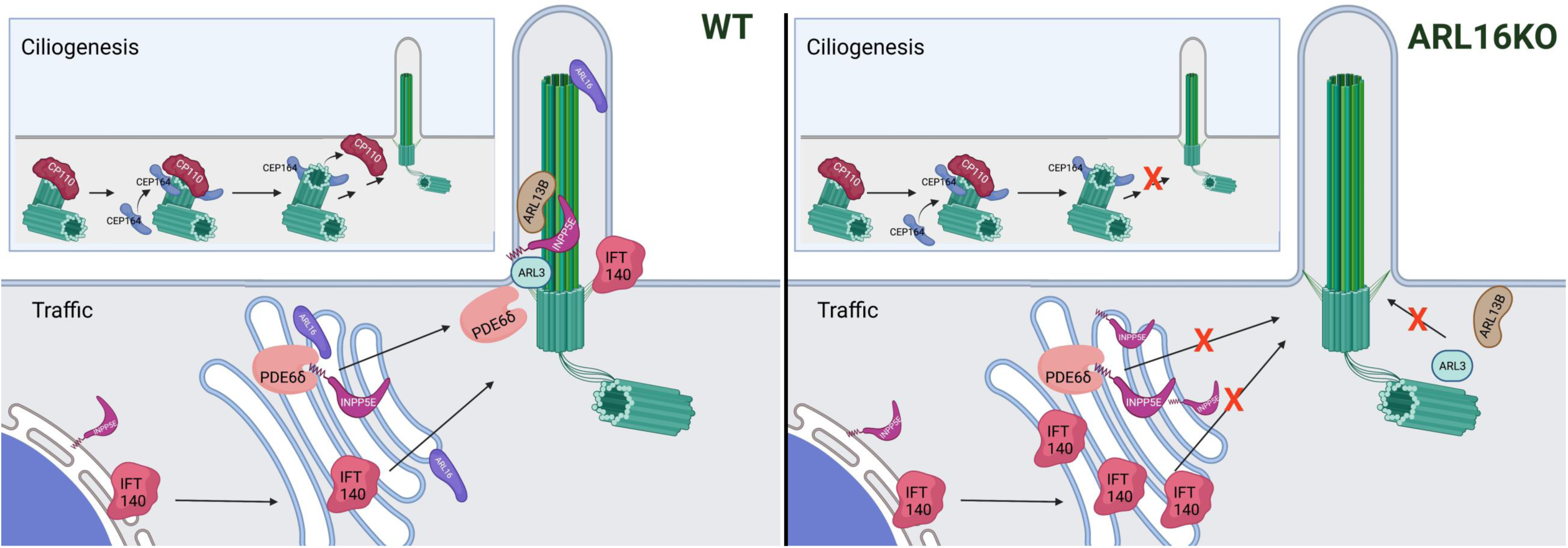
Model for ARL16 in ciliogenesis and Golgi to cilium traffic. *Ciliogenesis:* In WT cells induced to ciliate, CEP164 is recruited to mother centrioles as an early step in the ciliogenesis pathway, with later removal of the CP110 cap and extension of the axoneme. We propose that ARL16 acts after CP110 release to support ciliation such that in its absence ciliation is decreased and/or slowed. *Traffic:* In WT cells, INPP5E and IFT140 each traffic through the Golgi during transit to cilia. ARL16 supports the export of INPP5E from the Golgi, likely working with PDE6D, to traffic INPP5E from the Golgi to the cilium where it is released by ARL3 and retained by ARL13B. In contrast, IFT140 also requires ARL16, but not INPP5E, to be efficiently exported from the Golgi for transit to cilia. In addition, ARL3 and ARL13B fail to be recruited to or retained in cilia. Created with BioRender.com.

### Phylogenetic analyses predict role(s) for ARL16 in ciliary biology

Our phylogenetic analyses demonstrated that ARL16 is restricted to ciliated taxa, similar to the three ARL GTPases previously shown to have ciliary-related roles: ARL3, ARL6, and ARL13. The phylogenetic profile (or phyletic pattern) of a gene can be an extremely informative resource providing evolutionary as well as functional insights. Genes encoding proteins that underpin the cilium are a particularly good illustration of this concept. Numerous genes subsequently demonstrated to encode proteins representing constituents of cilia or being critical for its biogenesis had been originally predicted as candidate “ciliary genes” based on searching for genes whose phylogenetic distribution correlates with the ability of the taxa to form cilia (Avidor-Reiss et al., 2004; Li et al., 2004; Merchant et al., 2007; Nevers et al., 2017). Other ciliary genes were discovered by *ad hoc* observations, as is the case of the Ras superfamily protein RJL proposed to have a cilium-related role based on its phylogenetic distribution more than a decade ago (Elias and Archibald, 2009) and confirmed as a regulator of ciliogenesis recently (Piette et al., 2021).

ARL16 has been missed by most, if not all, previous comparative genomic screens for candidate ciliary taxa. One reason for this may stem from the high sequence divergence between ARL16 orthologs from distantly related eukaryotes, which might have resulted in the failure of commonly used, automated orthology prediction tools to recognize ARL16 genes in different species. Another confounding factor may be that the ARL16-cilium correlation is less conspicuous than is the case of many other ciliary genes in that it is missing in some ciliated species. That not all ciliary genes exhibit the same distribution pattern neatly correlating with the cilium is not unusual, as this cellular structure is highly malleable across species (Moran et al., 2014). Perhaps most obvious is the difference between motile and primary cilia as the latter (analyzed in this study) lack a central pair of microtubules, dynein arms and radial spokes. This phenomenon is documented also by our phylogenetically broad survey of the distribution of ARL3, ARL6, and ARL13 (Fig. 1; Table S1). None is present in all cilium-building species, with ARL3 being the least frequently missing. Hence, the absence of ARL16 from various ciliated eukaryotes by itself does not undermine the prediction that this GTPase has a specific cilium-associated role. In fact, the restriction of ARL3, ARL6, ARL13 and ARL16 to cilium-building eukaryotes can be interpreted as evidence that the cilium-associated role of these proteins, established by studying a limited set of model organisms, is a general property across eukaryotes as a whole. A prediction we can draw then is that the potential non-ciliary roles of these proteins are limited, making them dispensable whenever the cilium is lost. By extension, if ARL16 acts to regulate traffic from the Golgi, it may do so specifically in a Golgi-cilia pathway or for a subset of proteins destined for the cilium.

### Deletion of Arl16 causes a delay and overall decrease in ciliation yet increased ciliary length

We found that the percentage of cells that form cilia in response to serum starvation is reduced in *Arl16* KOs and that the affected step of ciliogenesis is downstream of CP110 uncapping of the basal body. It is by no means a complete defect in ciliation in the absence of ARL16. At later timepoints after serum starvation, *Arl16* KO cells achieve ciliation rates approaching WT MEFs, and by 72 hrs show no statistically significant differences from WT. More detailed tracking of centrioles and ciliation would be required to determine the precise step of ciliogenesis that is affected.

In addition, the cilia that do form in *Arl16* KOs are on average ∼90% longer than WT cilia (Fig. 3). While many things are known to result in shorter than average cilia (including the absence of ARL13B (Caspary et al., 2007), fewer things are known that increase ciliary length (Duran et al., 2016; Wang et al., 2021). One such regulator is Gli2-mediated autophagy of Ofd1 (Hsiao et al., 2018), though we did not explore this pathway in the *Arl16* KOs.

We recently discovered that ciliary rootlets are linked to ciliogenesis and the ARF GTPase activating protein (GAP) ELMOD2 (Turn et al., 2021). *Elmod2* KO resulted in rootlet fragmentation and increased centrosomal separation, each of which was reversed upon expression of fast cycling ARL2 (Turn et al., 2021). However, while the loss of ARL16 results in rootlet fragmentation, there is no difference in centrosome separation between *Arl16* KO and WT cells. These somewhat surprising results both suggest a role for ARL16 in rootlet integrity and may also raise questions about the role of rootlets in centrosome cohesion.

### ARL16 is required for localization of a subset of ciliary proteins

We found that ARL16 modulates multiple aspects of cilia. *Arl16* KO results in the marked reduction of several ciliary proteins including ARL13B, ARL3, and INPP5E. It is possible that the loss of some of these is a cascade effect beginning with the loss of ARL13B, as it has been shown to play important roles in binding and retention of both ARL3 and INPP5E in cilia (Gigante et al., 2020; Ismail et al., 2012; Kobayashi et al., 2009; Mourão et al., 2014; Schrick et al., 2006; Wiens et al., 2010). Farnesylated INPP5E is transported to the cilium by the carrier PDE6D where it is released by ARL3 (Fansa et al., 2016; Stephen et al., 2017). Therefore, the loss of INPP5E from cilia in *Arl16* KOs may result directly from the loss of ARL3 and/or ARL13B. We also found a reduction of AC3 in *Arl16* KO cilia, perhaps a result of altered ciliary PIP content due to the loss of INPP5E. Finally, changes in SMO recruitment (leading to defective Hh signaling) is likely due to the changes of these other ciliary proteins, each of which have been shown to regulate Hh signaling (Huangfu and Anderson, 2006; Humbert et al., 2012; Placzek and Briscoe, 2018).

### Golgi-cilia traffic is compromised by the lack of ARL16

The disruption of ciliary proteins often is accompanied by defects at the Golgi (Dafinger et al., 2011; Goncalves et al., 2010), and changes to the Golgi can affect ciliogenesis (Greer et al., 2014; Hurtado et al., 2011). While a few ciliary proteins also localize to the Golgi, (Baron Gaillard et al., 2011; Follit et al., 2006) the Golgi is close to the cilium both during ciliogenesis and after cilium maturation, suggesting that there is likely continuous transfer between these compartments (Evans et al., 2010; Poole et al., 1997). Some exclusive pathways transport specific cargos from the Golgi to the cilium, including rhodopsin and PKD2 (Kim et al., 2014; Ward et al., 2011). Furthermore, recent work has highlighted a Golgi to cilium traffic pathway for ciliary membrane proteins that is regulated by IFT-A (Quidwai et al., 2021).

IFT machinery is required for proper assembly and maintenance of primary cilia (Kannabiran, 2020; Lechtreck, 2015; Nachury, 2018; Roepman and Wolfrum, 2007). This machinery consists of two complexes; IFT-A, which is primarily responsible for retrograde transport from the cilium tip to the cell body, and IFT-B, which is critical for cilium assembly and anterograde transport. However, there is also mounting evidence of a role for IFT-A in anterograde transport as well (e.g., (Mukhopadhyay et al., 2010)). Both IFT-A and -B are multi-subunit complexes, with IFT-A containing 6 members (IFT140/144/139/122/121, and 43) and IFT-B further divided into two subcomplexes, a 10 subunit core (IFT-88/81/74/70/56/52/46/27/25, and 22), and a 6 subunit peripheral complex (IFT-172/80/57/54/38, and 20) (Jordan and Pigino, 2021). Little is known about where or how these complexes assemble in the cell prior to associating with cilia.

In *Arl16* KO cells, we found that the IFT-A core component IFT140 is lost from the cilium and accumulates at the Golgi. To date, the only other IFT protein known to localize to the Golgi is IFT20 (Follit et al., 2006), which is anchored there by binding to GMAP210 (Follit et al., 2008) and is required for opsin traffic from the Golgi to cilia (Keady et al., 2011). Loss of either IFT20 or IFT140 causes similar degenerative phenotypes in both the cilium and retinal photoreceptor cells, and loss of either abrogates ciliogenesis, though phenotypes are more severe in the IFT20 deletion (Crouse et al., 2014). However, several IFT subunits are associated with vesicular traffic functions in both ciliated and non-ciliated cells (Yang and Huang, 2019). Interestingly, our data do not show changes in other examined IFT-A or -B components, leading us to conclude that IFT140 accumulation at Golgi is independent of the other IFT components and that IFT140 may be involved in a novel Golgi to cilium traffic pathway, or perhaps traffic to a distinct compartment at which assembly of the IFT-A complex takes place.

In addition to IFT140, INPP5E accumulates at the Golgi of *Arl16* KOs. Traffic of INPP5E has been extensively studied as mutations in it are associated with both Joubert and MORM ciliopathies. Our data here build upon that model and add to it. We show that in both *Arl16* KOs and *Pde6d* KOs, INPP5E accumulates at the Golgi. High throughput screens also identified PDE6D as an interactor of ARL16 (Luck et al., 2020; Rolland et al., 2014). Therefore, we hypothesize that PDE6D picks up INPP5E at the Golgi to carry it to cilia in an ARL16-dependent manner.

### Summary

This initial analysis of ARL16 identified roles in Golgi-cilia traffic that are likely linked with its effects on the control of ciliary length and ciliation itself, though acting at distinct sites. Clearly, substantially more work is required to identify each of the sites and mechanisms by which ARL16 acts. The current study used mammalian cell culture and CRISPR/Cas9 introduced mutations as a model system, though the presence of ARL16 in almost all ciliated eukaryotes should provide alternative systems that are predicted to yield additional insights into ARL16 actions. Despite our survey of a limited number of ciliary proteins we identified a number of proteins whose presence in cilia is compromised by the loss of ARL16. These proteins have been shown previously to play important roles in Shh signaling (ARL13B, ARL3, INPP5E), axoneme integrity (ARL13B), phospholipid metabolism (INPP5E), and retrograde intraflagellar traffic (IFT140) acting at or in cilia. Our findings of INPP5E and IFT140 accumulation at the Golgi in *Arl16* KOs and INPP5E but not IFT140 at the Golgi in *Pde6d* KOs, support our model of ARL16 regulating two novel, independent traffic pathways from the Golgi to cilia. Finally, in a parallel study carried out in our lab we found phenotypes very similar to those reported here in *Arl16* KO lines when MEFs were deleted for either of the ARF/ARL GAPs ELMOD1 or ELMOD3 (Turn et al., 2021). In addition, expression of ARL16-myc in *Elmod1* or *Elmod3* KO cells was able to reverse those phenotypes, providing strong support for a model that includes these two proteins as acting in concert with ARL16 at the Golgi and in ciliary biology.

## MATERIALS AND METHODS

### Phylogenetic profiling of ARL3, ARL6, ARL13, and ARL16

The set of 114 eukaryotes previously analyzed for the composition of the ARF family gene complement (Vargova et al., 2021) was expanded by 25 additional eukaryotes (see Table S1), selected based upon the following criteria: (1) representatives of those major eukaryote lineages absent from the previous study (for the first time analyzing members of Hemimastigophora, Telonemia, Alveida, Rhodelphidia, and the CRUMs supergroup); (2) members of previously unsampled lineages known to have independently lost the cilium (Microsporidia, Myxozoa, Zygnematophyceae, Euglyphida); and (3) cilium-bearing close relatives of cilium-lacking taxa (e.g., the flagellated filasterean *Pigoraptor chileana* in contrast to the previously analyzed non-ciliated filasterean *Capsaspora owczarzaki*). Available genomic or transcriptomic data from these eukaryotes (sources provided in Table S2) were analyzed with BLAST searches (Altschul et al., 1997) to identify putative orthologs of ARL3, ARL6, ARL13, and ARL16. Sets of protein sequences predicted based on genomic assemblies and transcriptome assembly contigs were searched with BLASTP and TBLASTN, respectively, using an ARF1 sequence as a query. Because ARL16 is rather distant from other ARF family members and is not always identified with ARF1 as the probing query, we performed parallel analyses starting with a reference ARL16 sequence as the BLAST query. Hits with the e-value ≤0.1 were retrieved and blasted (with BLASTP and BLASTX in the case of protein and nucleotide sequences, respectively) against an in-house database of Ras superfamily protein sequences including the curated set of ARF family sequences reported previously (Vargova et al., 2021). Queries with the best 20 hits all corresponding to one of the four GTPases of interest (ARL3, ARL6, ARL13, and ARL16) were assigned as orthologs of the respective ARL protein, unless recognized as obvious contaminants from a different taxon (an issue sometimes happening in the case of transcriptome assemblies). Very rarely particular queries retrieved a set of 20 best hits consisting of a mixture of various ARF family members including some of the four focal GTPases. In all such cases the respective queries apparently corresponded to highly divergent genes with low sequence similarity to the canonical ARF family members, and were hence discarded. We cannot exclude the possibility that some of these genes as well as some of the unclassified ARF family members from the previous analysis (Vargova et al., 2021) are evolutionarily derived from ARL3, ARL6, ARL13, or ARL16 (*i*.*e*., being their orthologs or lineage-specific paralogs). However, in such cases their sequence divergence presumably entails a functional shift compared to canonical representatives of the four ARL paralogs, so failing to recognize their ultimate origin is unlikely to significantly impact the functional inferences from the phylogenetic distribution pattern of the four focal ARF family proteins. To avoid the possibility that any of the four GTPases is scored as missing for the given newly analyzed species only because of an accidental absence of the respective protein sequence resulting from inherently inaccurate genome annotation, we checked directly the genome assemblies with targeted TBLASTN searches and queries representing the ARL type not found in predicted protein sequence set. All newly identified ARL3, ARL6, ARL13, and ARL16 sequences were manually checked and if needed, the respective gene models were corrected to obtain accurate and complete protein sequences (the corrections are provided in Table S2).

The distribution pattern of ARL3, ARL6, ARL13, and ARL16 in the total set of 139 eukaryotes was correlated with the presence of cilia. Each species was scored as ciliated or non-ciliated (Fig. 1; Table S1) based on literature searches. A few apparently non-ciliated taxa (the foraminiferan *Reticulomyxa filosa* and the green algae *Ostreococcus lucimarinus* and *Coccomyxa subelipsoidea*) possess subsets of genes encoding characteristic ciliary proteins, suggesting that these species may form cilia at unknown life stages or lost the ability to build these structures only recently (Glöckner et al., 2014; Li et al., 2020). The cilia in these species, if indeed formed, are presumably highly reduced, owing to the lack of genes encoding many core ciliary proteins. Consistent with this, these species either completely lack ARL3, ARL6, ARL13, and ARL16 or, in the case of *R. filosa*, possess only one of these ARLs (ARL6) together with an apparent pseudogene corresponding to ARL3; Table S1). To account for the uncertain status of the cilium in these taxa, its presence was coded as ambiguous for the subsequent comparative analyses. The lack of evidence for the presence of ciliated stages in some poorly studied amoeboid protists (*Rigifila ramosa, Armaparvus languidus*) without genome sequence data should not be interpreted as evidence of absence of the cilium in these organisms, but the consistent lack of all four cilium-associated ARLs from the transcriptome data generated for them is consistent with the idea that all four, including ARL16, are not required when no cilium is formed. Hence, for the purpose of our correlation analysis, these species were scored as non-ciliated. Finally, the pelagophyte alga *Aureococcus anophagefferens* was not reported to have cilia, but it possesses three out of the four cilium-associated ARLs (Table S1) and was previously shown to encode various other ciliary proteins, so it likely forms a flagellated stage, most likely zoospores, like its relatives (for further details see Eliáš et al. (Eliáš et al., 2016)). Hence, we scored it as a ciliated eukaryote.

The strength of the dependence of the presence of ARL16 on the presence of the cilium was formally tested using the pairwise comparisons algorithm (Maddison, 2000) implemented in the Mesquite package (Maddison, W. P. and D.R. Maddison. 2021. Mesquite: a modular system for evolutionary analysis. Version 3.70. http://www.mesquiteproject.org). A strictly bifurcating tree representing the phylogenetic relationships of the 139 taxa analyzed was constructed on the basis of previously published molecular phylogenetic and phylogenomic analyses. The somewhat contentious branching order of the main eukaryote lineages was arbitrarily defined following the most recent and comprehensive phylogenomic study (Tice et al., 2021), but alternative topologies at the base of the eukaryote tree suggested by other studies would not impact the result of the correlation analysis, as they do not change the inferred positions of cilium and ARL16 loss events. The tree was loaded into Mesquite and a two-character matrix was built following the character coding presented in Table S1, with the character 1 corresponding to the cilium (three states: present, absent, and ambiguous) and the character 2 representing the presence or absence of ARL16. The Pairwise Comparison test was carried out with the “Most Pairs” option and 1000 pairings, and which gave the best tail p value range from 1.2⨯ 10^−7^ to 3.81⨯ 10^−6^ across all the pairings.

### Reagents, Antibodies, and Plasmids

All chemicals used were purchased from commercial sources. The following antibodies were used in these studies: ARL16 (1:100; Sigma; HPA043711), Acetylated Tubulin (1:1000; Sigma; T5192), Centrin clone 20H5 (1:1000; Sigma; 04-1624), polyclonal rabbit anti-centrin 3 (1:100 (Trojan et al., 2008)), myc (1:1000; Abcam; ab9132), HSP60 (1:1000; Stressgen; ADI-SPA-807), GM130 (1:1000; BD/Transduction; 610823), Tubulin (1:1000; EMD Millipore; MAB1864), β-COP (1:2,000; ThermoFisher PA1-061), FAPP2 (Gift from Antonella De Matteis), GBF1 (1:200; BD; 612116), CP110 (1:100; Proteintech; 66448-1-ig), gamma tubulin (1:1000; Sigma T6557 or Abcam ab11317), CEP164 (1:100; Santa Cruz; sc-515403), rootletin (1:1000; EMD Millipore; ABN1686), CEP290 (1:100; Proteintech; 22490-1-ap), NPHP4 (1:100; Proteintech; 13812-1-ap), IFT81(1:200; Proteintech; 11744-1AP), ARL13B (1:500; Proteintech; 17711-1-AP), ARL3(1:100; R75448 (Cavenagh et al., 1994), INPP5E (1:100; Proteintech; 17797-1-ap), AC3 (1:100; LSBio; LS-C204505/183274), SMO (gift from Kathryn Anderson (Ocbina et al., 2011), GLI3 (1:1000; R&D Systems; AF3690), IFT140 (1:200; Proteintech; 17460-1-AP), IFT144(1:100; Proteintech; 13647-1-AP), IFT88(1:200; Proteintech; 13967-1-ap), and IFT20(gift from Greg Pazour (Pazour et al., 2002)).

The plasmid directing expression of mouse ARL16-myc in mammalian cells was obtained by first having the open reading frame synthesized by GenScript and later using PCR to amplify this open reading frame with insertion of the C-terminal myc epitope (EQKLISEEDL) after a diglycine linker. The PCR product was ligated into pCDNA3.1 using KpnI and XhoI sites and the entire open reading frame was sequenced to confirm fidelity. The plasmid expressing SSTR3-GFP was a gift from Max Nachury (Marley et al., 2013).

### Cell culture

All cells were maintained in DMEM (ThermoFisher #11965) supplemented with 10% FBS at 37°C and 5% CO_2._ To induce ciliation, media was swapped with DMEM supplemented with 0.5% FBS for 48h unless otherwise indicated. Cells were grown in the absence of antibiotics. Routine screening for mycoplasma contamination was performed using DNA staining.

### CRISPR/Cas9

Genes of interest were disrupted in immortalized WT MEFs (ATCC CRL-2991) with CRISPR/Cas9 gene editing, as previously described (Schiavon et al., 2019; Turn et al., 2020; Turn et al., 2021). Guide RNA sequences targeting the coding region of the gene were designed using Benchling (benchling.com/academic). For *Arl16* the guides used were Guide 2: GGAGAGCCCCCACCGACGCGG and Guide 3: CGGAGATGGCAAAGGCGACCT. For *Pde6d* the guides used were Guide 1: GCAAATGGAAAAATTCCGCC and Guide 3: CACCGCCTTCGGGATGCCGAAACA.

Double stranded oligonucleotides of the guide sequences (with a substitution of a G for the first nucleotide to facilitate expression from the U6 promotor) were cloned into the pSpCas9(BB)-2A-Puro (PX459) V2.0 Vector (Addgene) at the BbsI sites. Cells were transfected with the resulting plasmid with a 1:3 ratio of DNA (2 µg) to Lipofectamine 2000 (6 µg) for 4 hrs in OptiMEM, according to manufacturer’s instructions. Cells were then replated and allowed to recover overnight in DMEM with 10% FBS. Cells were then grown in 3 µg/ml puromycin for 4 days, to enrich for transfected cells. Cells were then seeded into two 96-well plate at densities of 3-5 cells/well and monitored visually during growth. Clones resulting from single cells were isolated, expanded, and cryopreserved. PCR primers were designed to amplify genomic DNA outside the target site to allow sequencing of genomic DNA to identify and verify frameshifting mutations. Note that clones harboring no changes in the targeted region were often retained and referred to as “CRISPR WT” cells as they had been through the transfection, selection, and cloning process as the null lines and serve as another control against off target effects.

### Transfection

Cells were transfected with 4 µg DNA: 4 µL JetOPTIMUS transfection reagent (VWR, 76299-634) according to manufacturer guidelines in standard medium overnight. The next day, cells were replated on coverslips in standard medium and allowed to recover for 24 hr. Cells were then serum starved for 24-72 hr to induce ciliation prior to fixation.

### Immunofluorescence

All cells were cultured on coverslips coated with matrigel (BD Biosciences #356231).

#### PFA protocol

For the following antibodies (Ac-Tu, ARL13B, ARL3, INPP5E, γ-tubulin, SMO, AC3), cells were fixed 4% paraformaldehyde (PFA) in PBS pre-warmed to 37°C on the benchtop for 15 min. Cells were permeabilized with 0.1% Triton-X 100 in PBS for 10 minutes at room temperature. For Ac-Tu, ARL13B, and SMO, cells were blocked with 1% bovine serum albumin (Sigma #A3059) in PBS for 1 hr. Primary antibodies were diluted in blocking solution and applied to cells at 4° C overnight. Cells were washed 4 × 5 min with PBS before incubating with secondary antibodies (1:1000 Alexa fluorophores, ThermoFisher) in blocking solution for 1 hr at room temperature. Cells were washed 4 × 5 min with PBS before being mounted onto slides with MOWIOL. For ARL3, INPP5E, and AC3, a blocking solution of 10% FBS in PBS was used in place of 1% BSA.

#### Methanol protocol

For these antibodies (γ-tubulin, Ac-Tu, INPP5E at Golgi, CEP164, and CEP290), cells were fixed with methanol at -20º C for 10 min, washed in PBS with agitation. Cells were blocked with blocking buffer (PBS with 10% FBS) for 30 min at room temperature. Primary antibodies were diluted in blocking buffer and applied to the cells at 4° C overnight. Cells were washed 3 × 5 min with PBS with agitation before applying secondary antibody diluted 1:500 in blocking buffer for 1 hr at room temperature. Cells were then washed and mounted as described above.

#### IFT protocol

For IFT staining (IFT140, IFT144, IFT88, and IFT81). Immediately out of the incubator, cells were washed 2 x in PBS warmed to 37° C prior to fixation with 4% PFA in PHEM (60 mM PIPES, 22 mM HEPES, 10 mM EGTA, 4 mM MgSO_4_-7H_2_O, pH 6.9) for 15 min at room temperature. Cells were washed 2x with PBS and treated with 50 mM NH_4_Cl twice for 15 min. Cells were washed again with PBS before permeabilization with 0.25% Triton-X100 in PBS for 10 min at room temperature. Cells were blocked with 10% FBS in PBS with 0.2% Tween-20 for 60 min at room temperature and then incubated with primary antibodies in 1% FBS in PBS with 0.025% Triton-X100 at room temp for 1 hr or at 4° C overnight. Cells were washed 4×10min with PBS before incubation with secondary antibodies for 60 min at room temp. Cells were rinsed with 0.25% Triton-X100 in PBS 5×10 min before mounting as above.

### Microscopy

Samples were visualized using an Olympus IX81 widefield microscope and Slidebook software; 100X magnification [UPIanFI, 1.30 NA Oil]. Images were processed and analyzed using FIJI image analysis software. Any images appearing in the same panel of a figure were processed identically including objectives, acquisition settings, cropping, brightness adjustments, and any other processing settings.

### Scoring of cell phenotypes

For all phenotypes that were scored, experiments were performed in triplicate and scored at least in duplicate as indicated in the corresponding figure legends. For percent ciliated cells, cilia were identified using Ac-Tu. For presence of markers in cilia (ARL13B, ARL3, INPP5E, etc.), we binned them as either present (visible even without checking the Ac-Tu channel), reduced (present, but only noticeable upon switching to Ac-Tu channel), or absent (cannot be detected upon switching to Ac-Tu channel). For all ciliary phenotyping, Ac-Tu was used to define cilia. For centrosomal/basal body scoring, γ-tubulin was used as the standard comparison point. Finally, for Golgi staining/localization, GM130 was used to define the Golgi.

### Human Tissue

The human donor eye tissue applied in the present study was obtained 11.5 hr post mortem from a female donor (# 252-09), 65 years of age without any underlying health conditions, from the Department of Ophthalmology, University Medical Center Mainz, Germany. The guidelines to the declaration of Helsinki (http://www.wma.net/en/30publications/10policies/b3/) were followed.

### Immunohistochemistry of Retinal Sections

Human retinae were dissected from enucleated eye balls and cryofixed in melting isopentane and cryosectioned at -20° C in a cryostat (HM 560 Cryo-Star; MICROM) as previously described (Karlstetter et al., 2014; Wolfrum, 1991). 10 µm sections were placed on poly-L-lysine-precoated coverslips and incubated with 0.01% Tween 20 in PBS for 20 min. After washing, sections were flooded with blocking solution (0.5% cold-water fish gelatin plus 0.1% ovalbumin in PBS) and incubated for at least 30 min followed by an overnight incubation with primary antibodies at 4° C in blocking solution (Trojan et al., 2008). Washed cryosections were incubated with secondary antibodies conjugated to Alexa 488 or Alexa 568 (Invitrogen) in blocking solution and with DAPI (Sigma-Aldrich) to stain the DNA of nuclei, for 1.5 h at room temperature in the dark. After three washes in PBS, specimens were mounted in Mowiol 4.88 (Hoechst) and imaged using a Leica DM6000B deconvolution microscope.

### Shh assay

Shh response was determined by measuring transcriptional changes in Gli1 and Ptch1 mRNA levels, as previously described (Mariani et al., 2016). Cells were maintained in low serum (0.5% FBS) media for 48 hrs either with or without Shh conditioning. RNA was prepared using the Qiagen RNeasy Kit with QIAshredder homogenizer columns, according to the manufacturer’s protocols. RNA (200 ng) was used to generate cDNAs using BioRad iScript Reverse Transcription supermix. The following primers were used for quantitative PCR (qPCR):

#### Pold3 (housekeeping gene)

F: 5′-ACGCTTGACAGGAGGGGGCT-3′

R: 5′-AGGAGAAAAGCAGGGGCAAGCG-3′

#### Gli1

F: 5′-CTTCACCCTGCCATGAAACT-3′

R: 5′-TCCAGCTGAGTGTTGTCCAG-3′

#### Ptch1

F: 5’-TGCTGTGCCTGTGGTCATCCTGATT-3’

R: 5’-CAGAGCGAGCATAGCCCTGTGGTTC-3’

In brief, the cDNA was combined with primers and Bio-Rad SsoAdvanced Universal SYBR Supermix according to the manufacturer’s protocols (1725270). Samples were run on a Bio-Rad CFX96 Touch Real-Time PCR Detection System, and data were analyzed using Bio-Rad CFX Manager 3.1. The following program conditions were used: 95° C for 5 min; 45 cycles of 95° C for 15 s; 57 °C for 30 s. Reactions were performed in technical duplicate on three biological replicates. Data were then analyzed by the ΔΔCT method and normalized to control WT levels for each transcriptional target (Livak and Schmittgen, 2001).

### Reproducibility and Statistics

All data were plotted using GraphPad Prism. Error bars represent the standard error of the mean (SEM), and box-and-whisker plots indicate the range of the data along with the median and upper/lower quartiles. T-tests or one way ANOVA tests were used to determine whether there were significant differences between test groups as indicated. Asterisks in a figure indicates statistical significance: *p < 0.05, **p < 0.01, ***p<0.001. Actual p values are indicated in the figure legends.

## Abbreviations include

Ac-Tu: acetylated tubulin
GAP: GTPase activating protein
GEF: guanine nucleotide exchange factor
KO: knock out cell line
MEF: mouse embryonic fibroblast.

## ACKNOWLEDGEMENTS

This work was supported by grants from the NIH (R35GM122568 to RAK, R35GM122549 to TC, and F31HD096815 to SID), ERD Funds, project OPVVV CZ.02.1.01/0.0/0.0/16_019/0000759 (Centre for research of pathogenicity and virulence of parasites), Czech Science Foundation grant 20-27648S and ERD Funds, project OPVVV CZ.02.1.01/0.0/0.0/16_019/0000759 (to M.E.). We also thank Mike Murphy, RefSeq Curator (NCBI/NLM/NIH) for clarification on mRNA variants of the human ARL16 mRNA. The authors regret that due to the journal’s limitation on words in the text we are unable to cite many important contributions in the primary literature.

## FIGURE LEGENDS

**Figure S1:**
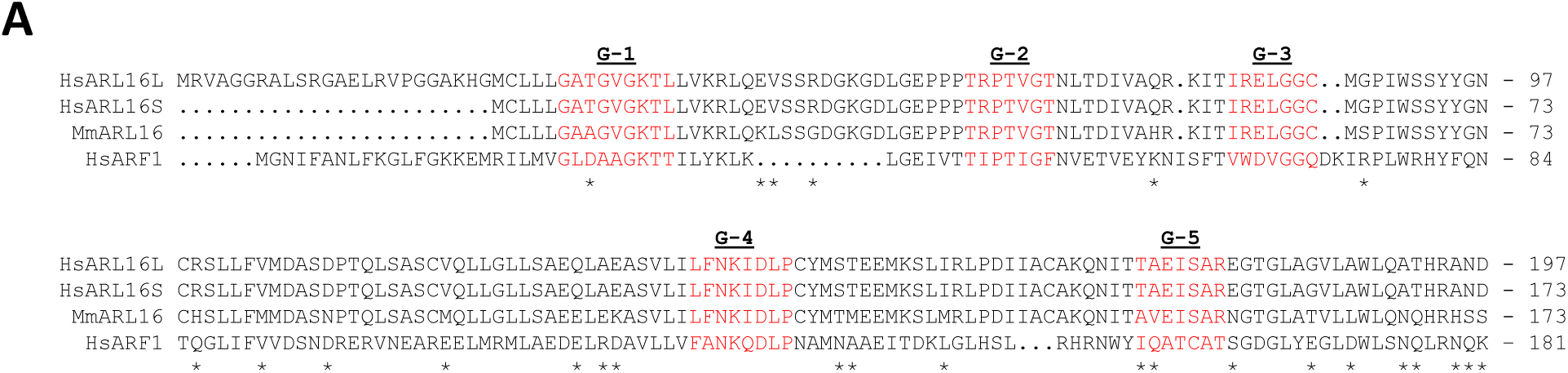
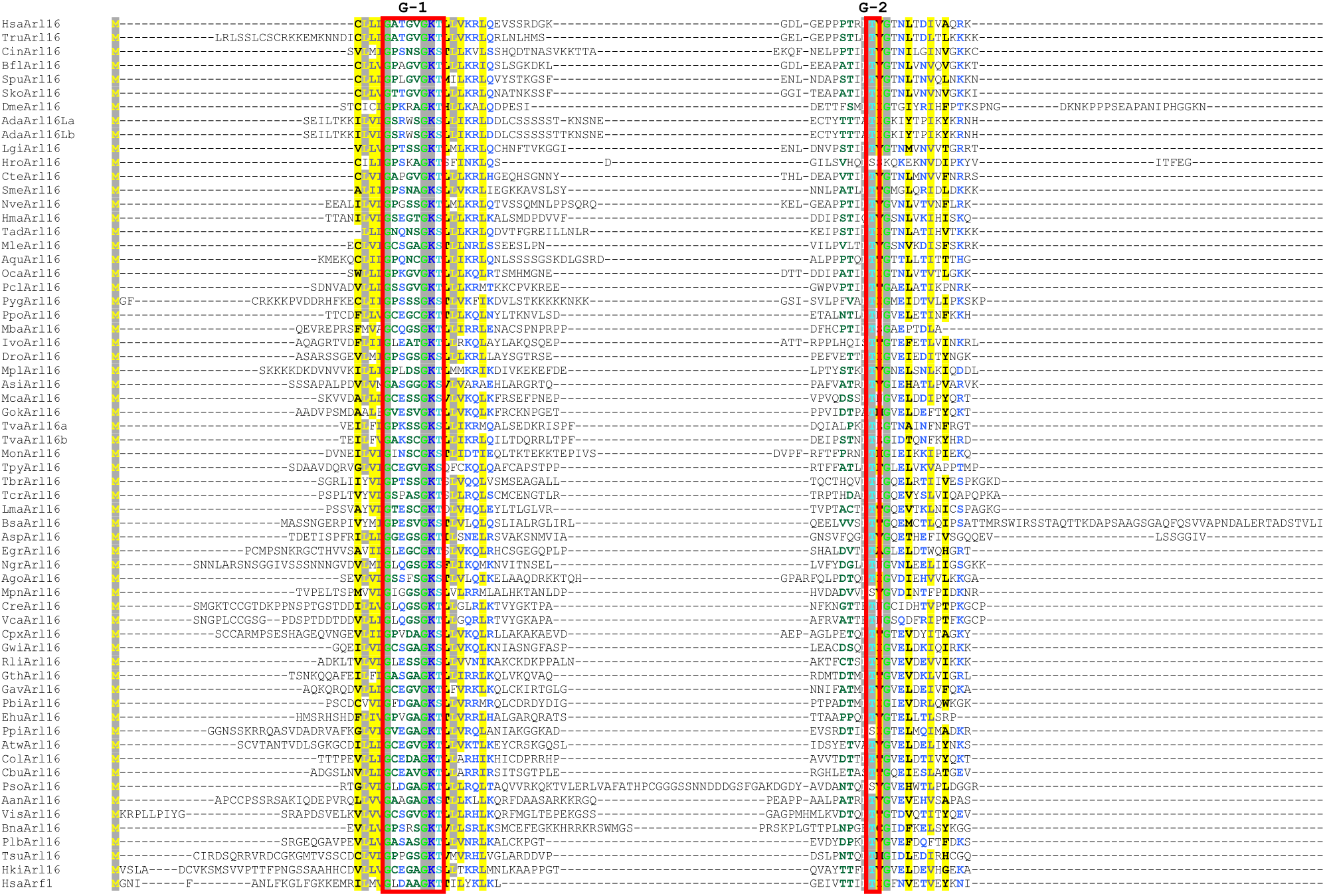
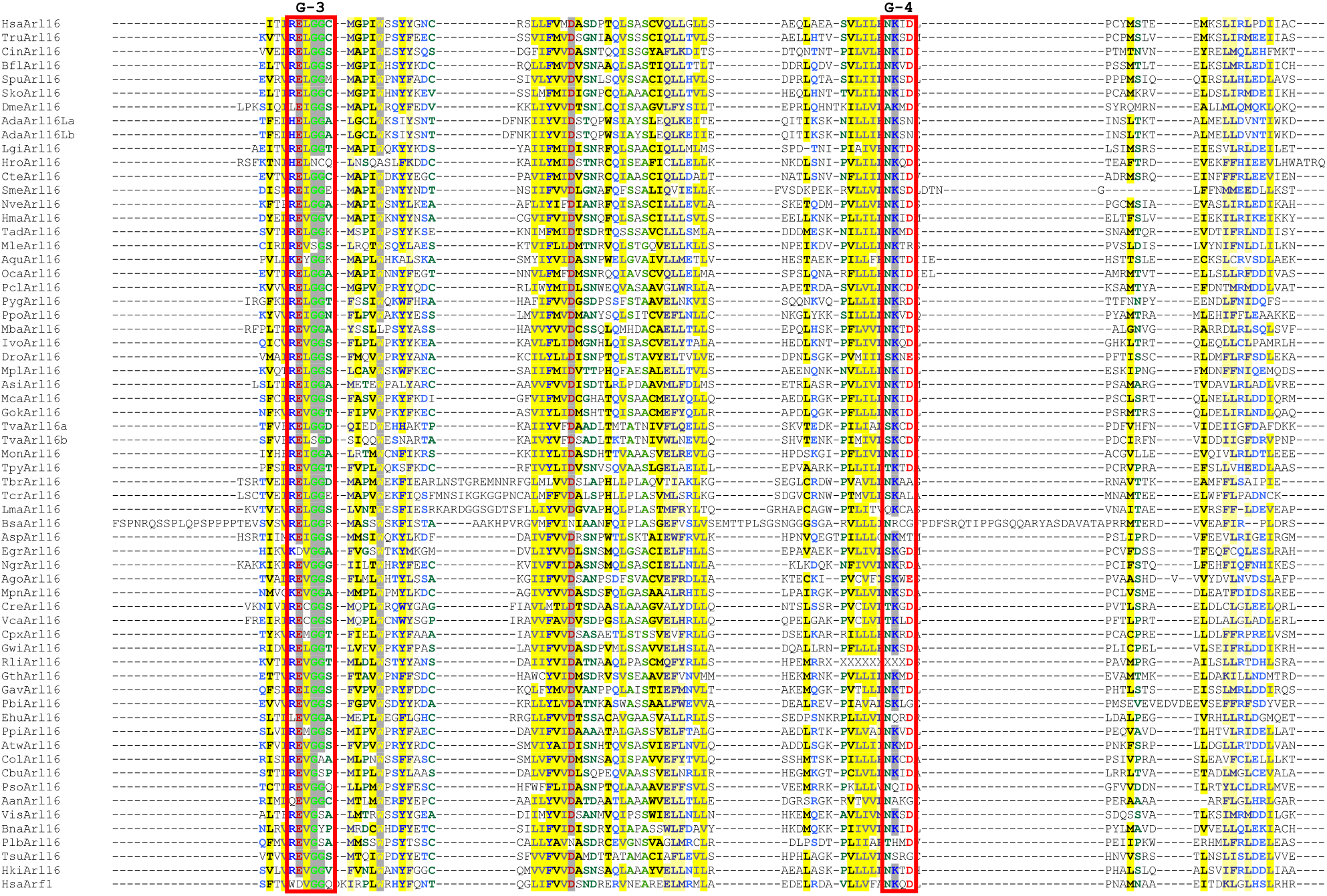
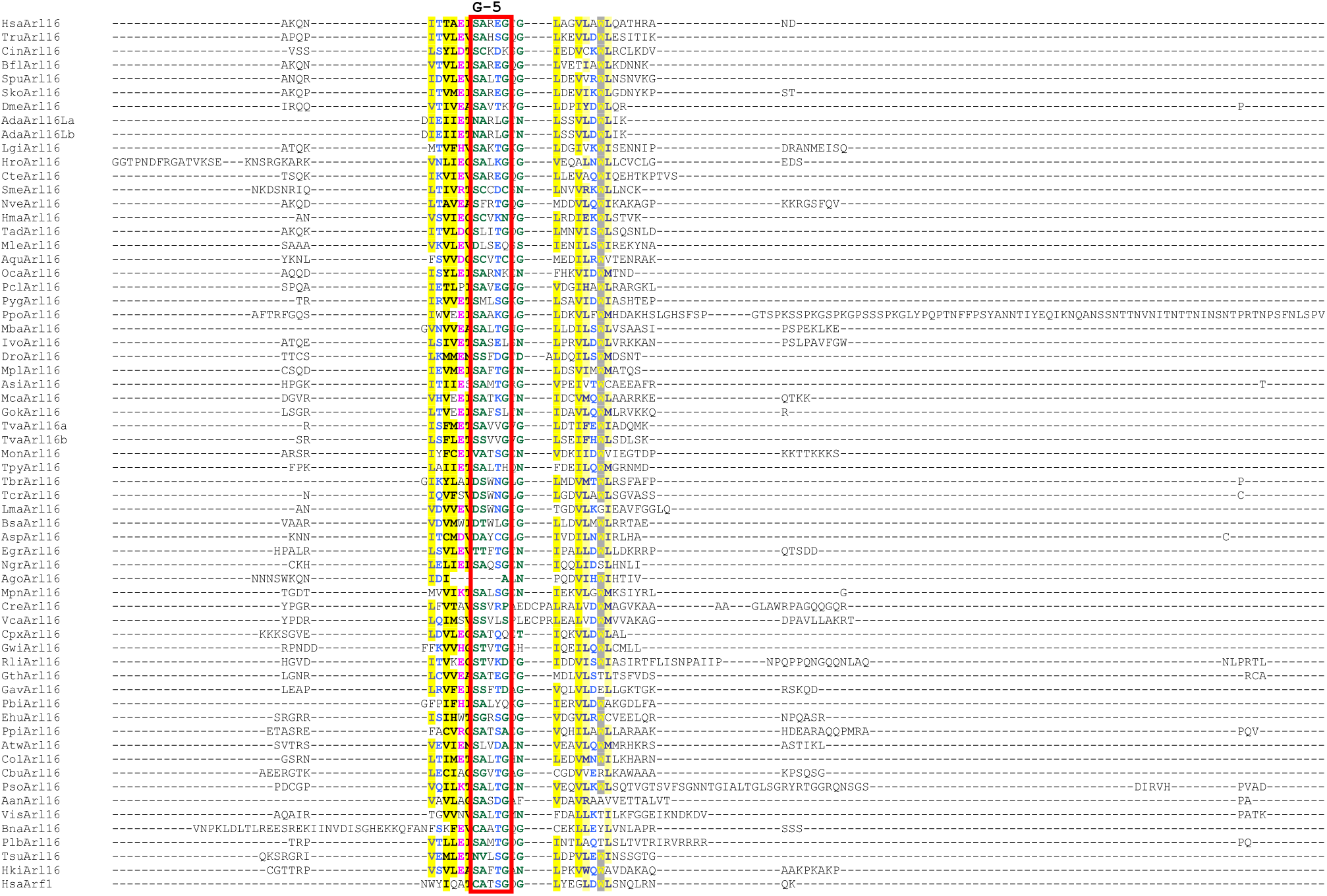
ARL16 protein sequences diverge from ARF1 in key motif residues. **(A)** Alignment of mouse and human ARL16 and human ARF1 protein sequences showing both the long and short human forms and the human ortholog. G motifs are highlighted in red. The mouse and short human forms contain a Cysteine residue at position 2, which may be palmitoylated. Asterisks (*) indicate residues that differ between human (short) and mouse ARL16 proteins. Alignment was performed using Multalin (http://multalin.toulouse.inra.fr/multalin/multalin.html). **(B)** Multiple alignment of ARL16 protein sequences from across the eukaryote phylogeny. All identified ARL16 orthologs from the sample of 139 phylogenetically diverse eukaryotes analyzed in this study and (for comparison) the human ARF1 sequence (HsaArf1) were aligned using MAFFT version 7 (https://mafft.cbrc.jp/alignment/server/) and residue conservation pattern was visualized by coloring the alignment using CHROMA Version 1.0 (http://www.lg.ndirect.co.uk/chroma). Four characteristic G-motifs are framed by red boxes. Note that the ARL16 protein from *Physarum polycephalum* (PpoArl16) has a very long C-terminal extension consisting of low-complexity repeats; this part of the protein was omitted from the figure. Sequence labels include three-letter species abbreviations indicated in Table S1. Sources of the sequences are provided in Table S2. All the sequences included were checked for accuracy of their inference from nucleotide sequence data, so the apparently unusual features of some of them (insertions, deletions or poor conservation of some regions) most likely reflect the actual primary structure of the respective proteins. In a few cases there is uncertainty concerning the N-terminus of the proteins, as it is possible some (*e*.*g*., TruArl16) are artificially extended upstream of the real initiating (N-terminal) methionine residue. The human sequence (HsaArl16) is represented by the shorter variant.

**Figure S2:**
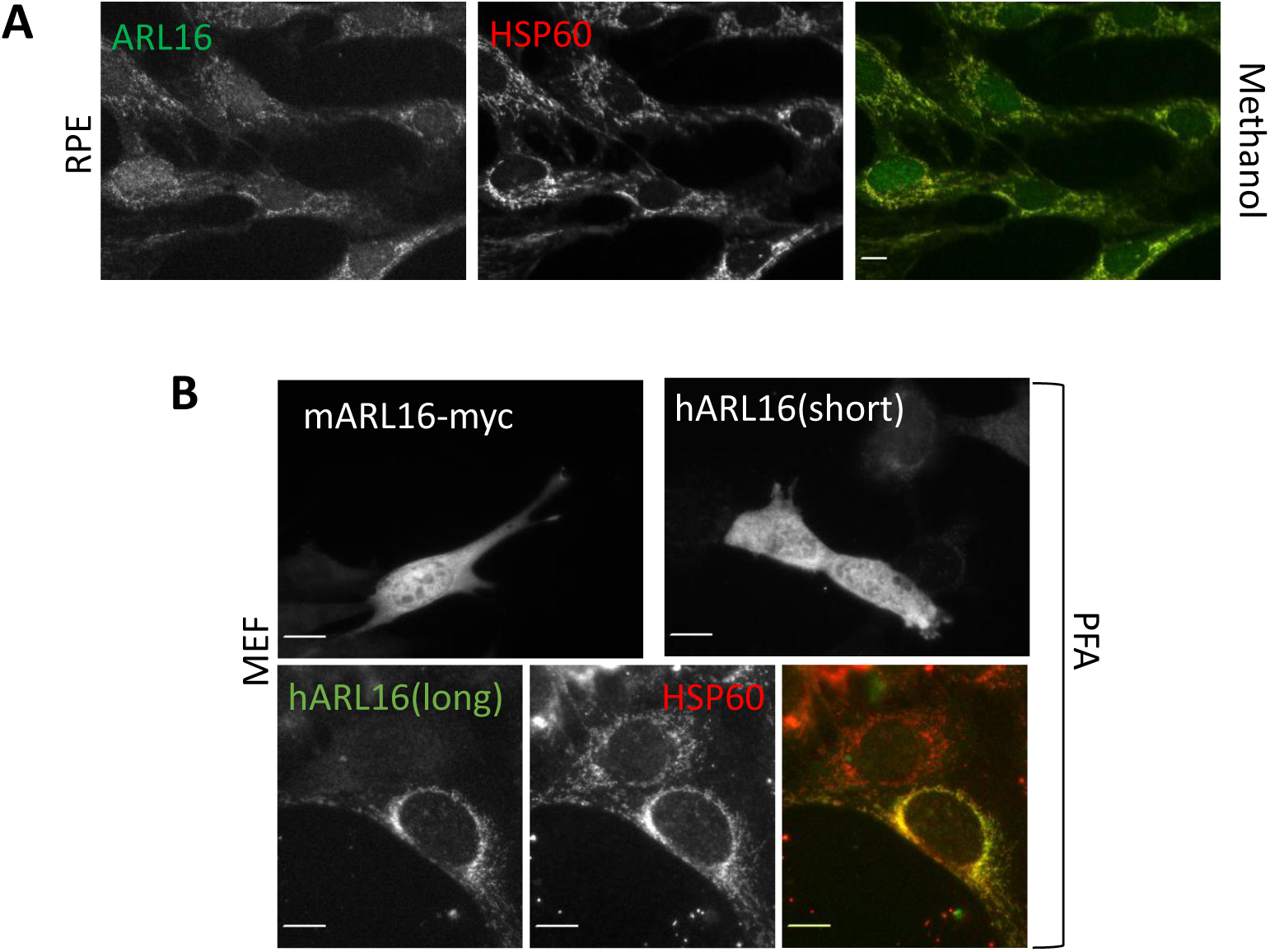
ARL16 localizes to cytosol and mitochondria. **(A)** Endogenous ARL16 can be detected at mitochondria in RPE1 cells after methanol fixation, as indicated by colocalization with HSP60. Cells were serum starved for 24 hrs prior to fixation for 5 mins in cold methanol, as described under Materials and Methods. Cells were stained for HSP60 (red) and ARL16 (green). Scale bar = 10 µm (60x). **(B)** Exogenously expressed mouse ARL16-myc and human ARL16 (short) localize primarily to cytosol (diffuse staining) in WT MEFs. Cells were transfected, serum starved for 24 hr, fixed with PFA as described in *Materials and Methods*, then stained for myc or ARL16. Exogenously expressed human ARL16(long) localizes to mitochondria when expressed in MEFs, as indicated by co-localization with HSP60 (bottom panels). Scale bar = 10 µm. (100x).

**Figure S3.**
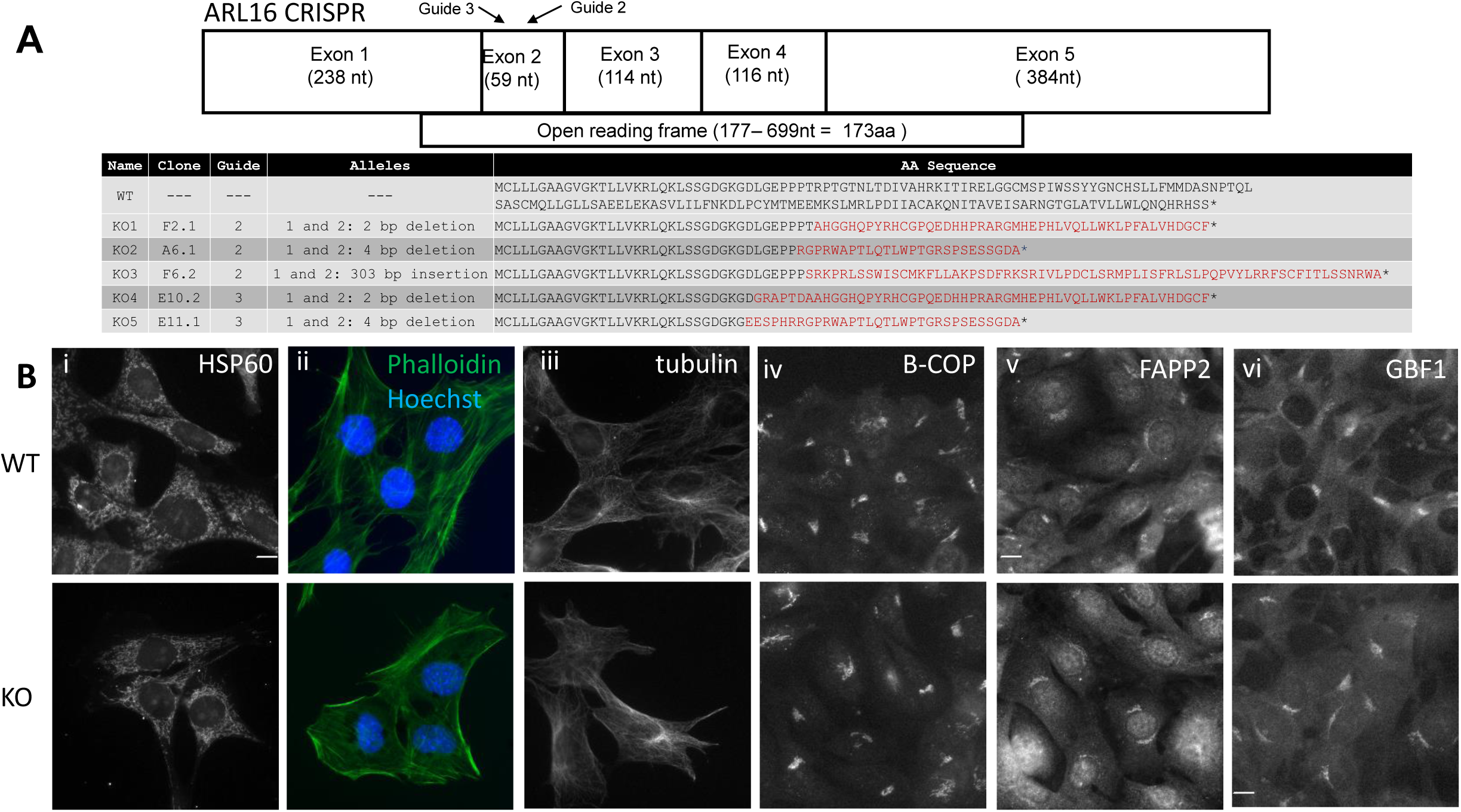
Knock out of *Arl16* in MEFs and screening of markers. (**A**) Schematic of mouse *Arl16* transcript, with the open reading frame shown below it. Arrows indicate sites targeted by the guide RNAs used for genome editing. Summary of alleles of the 5 *Arl16* KO lines. We obtained three clones from guide 2 and two clones from guide 3. Each of the clones have frameshifting mutations that lead to premature termination of the protein sequence. The black sequence indicates the WT amino acid sequence. Red sequence indicates the predicted amino acids following the frameshift mutations. **(B)** Mitochondria, actin, microtubules, and nuclei are unchanged in *Arl16* KOs. (i) WT and *Arl16* KO cells (KO3) were fixed with PFA and stained for HSP60 to mark mitochondria, (ii) with phalloidin (green) to mark the actin cytoskeleton and Hoechst (blue) to mark nuclei, (iii) for tubulin to mark the microtubule cytoskeleton, and (iv-vi) β-COP, FAPP2, and GBF1 to mark the Golgi. Scale bar=10 µm (60x).

**Figure S4.**
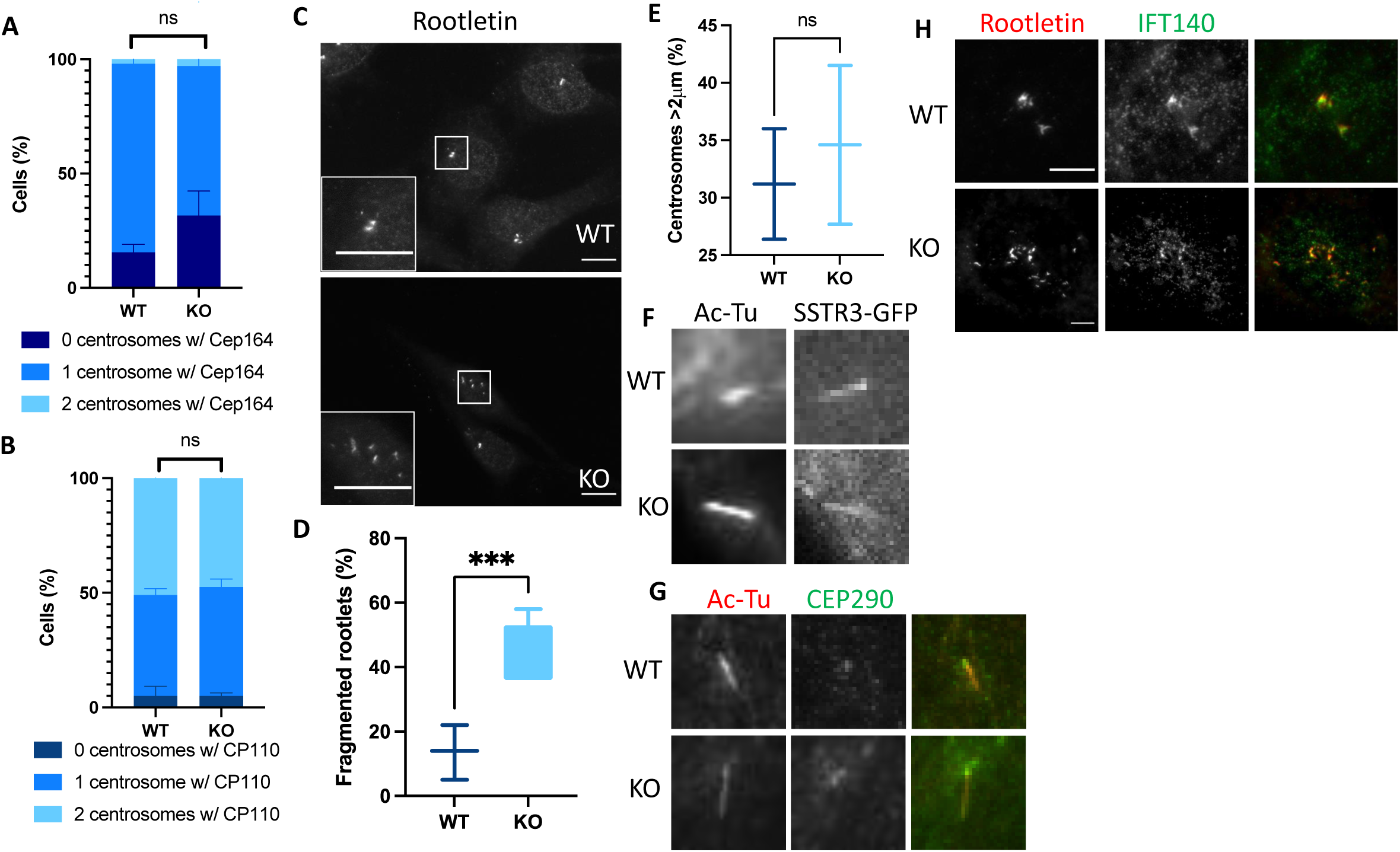
A number of ciliary features are unchanged between WT and *Arl16* KO lines. *Arl16* KO cells displayed levels of (**A**) CEP164 as well as (**B**) CP110 staining at centrioles comparable to WT MEFs. Cells were serum starved for 24 hr and stained for γ-tubulin and CEP164 or CP110. The percentage of cells with CEP164 or CP110 at centrosomes was counted. N=2 × 100 cells x 1 line for WT or 2 lines for *Arl16* KOs. **(C)** Rootlets are fragmented in *Arl16* KO cells. WT and *Arl16* KO cells were serum starved for 24 hr and stained for rootletin. Scale bar = 10 µm (100x). **(D)** Quantification of C. N= 2 × 100 cells for 1 WT line and 4 *Arl16* KO lines. (**E**) There is no difference in centrosome separation between WT and *Arl16* KO cells. Cells were serum starved for 24 hr and stained for γ-tubulin. The distance between γ-tubulin positive puncta (centrosomes) was measured using FIJI. (**F**) SSTR3-GFP reaches cilia indistinguishably in *Arl16* KO and WT cells. Cells were transfected with plasmid directing expression of SSTR3-GFP, the next day replated and allowed to recover for 24h followed by 24h of serum starvation prior to fixation and staining for Ac-Tu. (**G**) CEP290 localizes normally to the transition zone of *Arl16* KO cells. Cells were serum starved for 24h and stained for Ac-Tu and CEP290. (**H**) IFT140 localizes to rootlets and rootlet fragments in WT and *Arl16* KO cells. Cells were serum starved for 24h, fixed with methanol, and stained for rootletin and IFT140, Scale bar = 10 µm (100x).

**Figure S5:**
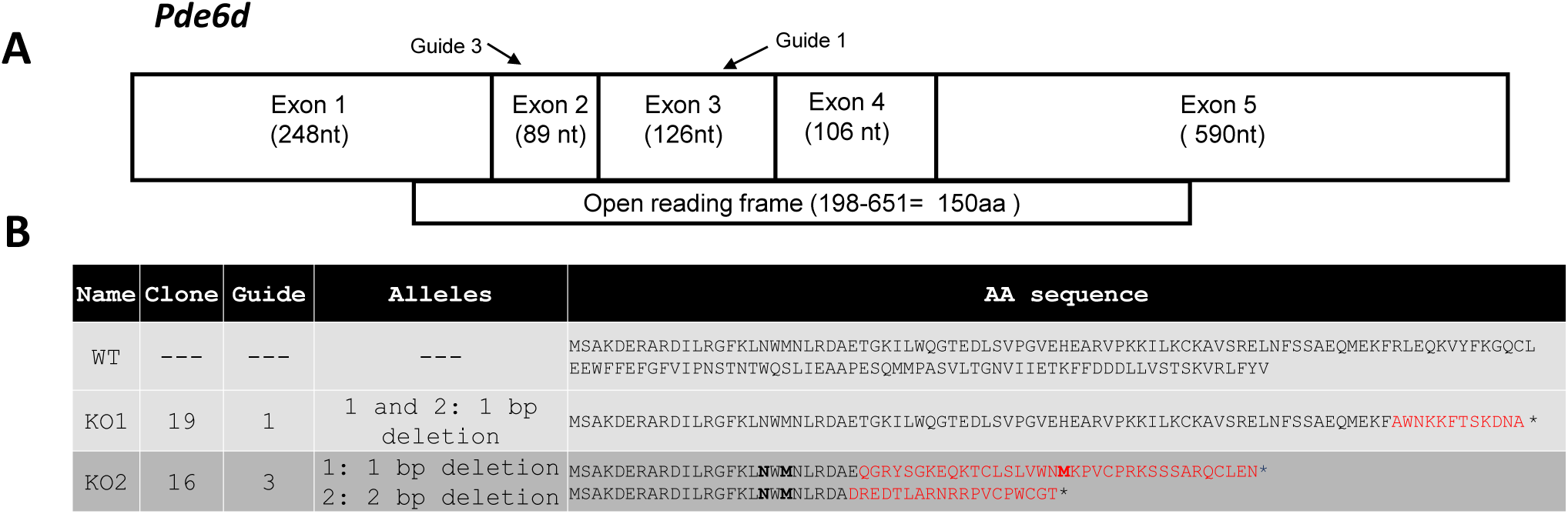
*Pde6d* CRISPR design. (**A**) Schematic of mouse *Pde6d* transcript, with the open reading frame shown below it. Arrows indicate sites targeted by the guide RNAs used for genome editing. (**B**) Summary of alleles of the 2 *Pde6d* lines used in these studies. Each of the clones have frameshifting mutations that lead to premature termination of the protein sequence. The black sequence indicates the WT amino acid sequence. Red sequence indicates the predicted amino acids following the frameshift mutation.

## Notes

### Competing Interest Statement

The authors have declared no competing interest.

